# Variable ectopic heterochromatin islands provide an alternative route to antifungal heteroresistance in *Cryptococcus neoformans*

**DOI:** 10.64898/2026.06.10.731255

**Authors:** Rebecca Yeboah, Sandra Catania, Pin Tong, Maria Isabel Navarro-Mendoza, Carlos Perez-Arques, Manu Shukla, Alison L. Pidoux, Neil R.H. Stone, Tihana Bicanic, Joseph Heitman, Robin C. Allshire

**Affiliations:** Centre for Cell Biology and Institute of Cell Biology, School of Biological Sciences, University of Edinburgh. Edinburgh EH9 3BF, Scotland, UK; Institute of Infection and Immunity, School of Health and Medical Sciences, City St. George’s University of London, London SW17 0RE, UK; Department of Molecular Genetics and Microbiology, Duke University School of Medicine, Durham, North Carolina 27710, USA

**Keywords:** Chromatin, Epigenetics, Epimutation, Fluconazole, Aneuploidy, Antimicrobial Resistance, Antifungal, Unstable, Histone, Adaptation

## Abstract

The fungal pathogen *Cryptococcus neoformans* is estimated to cause approximately 180,000 deaths annually, primarily among immunocompromised HIV patients in resource-limited countries in central and southern Africa. Economic constraints regularly restrict treatment to fluconazole monotherapy. Unstably resistant cells arise in any otherwise fluconazole-sensitive *C. neoformans* population, with such ‘heteroresistance’ often involving chromosome 1 disomy. Here we uncover a form of heteroresistance that occurs by a mechanism distinct from aneuploidy. Transient islands of histone H3 lysine 9 methylation-dependent heterochromatin occur at several locations across the genomes of clinical isolates from African patients. Some islands dissipate in the absence of fluconazole selection but are restored upon re-exposure. Deletion of specific heterochromatin island-located genes increased fluconazole resistance in an otherwise wild-type background, suggesting that heterochromatin-mediated alterations in gene expression elicit unstable resistance. Thus, in addition to aneuploidy, transient heterochromatin islands also appear to contribute to *C. neoformans* fluconazole heteroresistance. Heterochromatin islands have recently been shown to confer unstable antifungal resistance in the ascomycete fungus *Schizosaccharomyces pombe* (fission yeast) and fungal species of the *Mucor circinelloides* complex. Thus, similar unstable resistant epimutations may be widespread in pathogenic fungi, however, their instability and inaccessibility to detection by direct sequencing poses challenges for monitoring this form of heteroresistance in clinical settings.

## Introduction

Invasive pathogenic fungi are responsible for approximately 2.5 million human deaths per annum (Denning 2024). Immunocompromised individuals are particularly susceptible to invasive fungal infections (Loyse et al. 2019). Furthermore, diagnosis and treatment are hampered by the lack of inexpensive, accurate tests and effective antifungals in resource-limited settings (Loyse et al. 2019; Hurt et al. 2021). *Cryptococcus neoformans* has been designated by the World Health Organisation as one of the four critical priority human fungal pathogens that pose the greatest public health threat (Fisher and Denning 2023).

*C. neoformans* is an environmental saprophyte that enters human hosts by inhalation (May et al. 2016; Hallas-Møller et al. 2024). Initially infecting the lungs, it can eventually spread to the bloodstream and central nervous system (CNS) (May et al. 2016; Bednarek and Brown 2024). In healthy individuals, infections are generally innocuous and either cleared from the lungs or enter a dormant or latent state; however, the development of CNS cryptococcosis in HIV-infected patients results in 100% mortality if not treated, with a mortality rate of up to 50% even when treated. The estimated total number of deaths worldwide due to *C. neoformans* infections is approximately 180,000 per annum, with most cases occurring in sub-Saharan Africa (May et al. 2016; Loyse et al. 2019; Hurt et al. 2021)

The number of antifungals available for treating patients with cryptococcosis is limited to three therapies: the azole fluconazole (FLC), the polyene amphotericin B (AmB) and the pyrimidine analogue flucytosine (5-FC). Initial treatment for cryptococcosis involves combination therapy using AmB combined with 5-FC, followed by consolidation and prolonged maintenance with FLC (Loyse et al. 2019; Hurt et al. 2021). Unfortunately, despite advocacy efforts, access to AmB and 5-FC treatments remains limited in sub-Saharan Africa. Thus, oral FLC monotherapy remains the most widely available treatment for cryptococcal infections (Loyse et al. 2019; Hope et al. 2019; Hurt et al. 2021).

The drawback of FLC monotherapy is that *C. neoformans* exhibits a phenomenon known as heteroresistance, whereby a fraction of cells (<1-10%) in any population exhibits transient FLC resistance that is readily lost in the absence of drug selection (Sionov et al. 2009; Hope et al. 2019). Heteroresistance to FLC in *C. neoformans* has been described as intrinsic adaptive resistance because such resistant subpopulations are universal, even in isolates not previously exposed to FLC, and is selected for upon drug treatment. Resistance returns to initial background levels following drug removal (Sionov et al. 2009; Hope et al. 2019).

Heteroresistance in *C. neoformans* is usually attributed to aneuploidy, most frequently resulting from chromosome 1 disomy (Sionov et al. 2009; Hope et al. 2019; Sionov et al. 2010). FLC targets the Erg11/Cyp51 lanosterol 14α-demethylase, a cytochrome P450 enzyme required for ergosterol synthesis and the integrity of the plasma membrane and other membranes (Sagatova et al. 2015; Sionov et al. 2013). The presence of an additional copy of chromosome 1 doubles dosage of the genes encoding Erg11 and the Afr1 efflux pump, which reduces intracellular levels of FLC. In the absence of drug selective pressure, loss of the chromosome 1 disomy occurs through chromosome nondisjunction, and the resulting 1N euploid state has a growth advantage compared to the aneuploid state, restoring drug sensitivity. Consequently, chromosome 1 disomy confers FLC heteroresistance to only a fraction of cells in any *C. neoformans* population, even in the absence of previous FLC selection. Intriguingly, however, an unexplained form of FLC heteroresistance was observed in a subset of *C. neoformans* samples collected directly from the cerebrospinal fluid of a cohort Tanzanian HIV/AIDS patients with cryptococcal meningitis (Stone et al. 2019). All 20 samples were plated directly on FLC-containing plates to capture heteroresistant subpopulations, but eight of the samples that exhibited FLC heteroresistance were found to be euploid (i.e. non-aneuploid).

Transient adaptive phenotypes, such as drug resistance, can also result from the formation of RNAi-dependent or chromatin-mediated epimutations in fungi, plants and animals (Heard and Martienssen 2014; Miska and Ferguson-Smith 2016; Ganem and Sarkies 2026). The pathogenic fungus *Mucor circinelloides* species complex has been shown to develop epimutants which exhibit reversible unstable resistance to the antifungal compounds FK506 and rapamycin through RNAi-dependent or heterochromatin-mediated mechanisms (Calo et al. 2014, 2017; Navarro-Mendoza et al. 2023; Son et al. 2026). Although transient, these epimutant states can be transmitted through sexual reproduction (Pérez-Arques et al. 2025). In wild-type fission yeast, *Schizosaccharomyces pombe*, histone H3 lysine 9 methylation (H3K9me)-dependent heterochromatin can arise at novel chromosomal locations that are distinct from the constitutive domains of heterochromatin at centromeres, telomeres and the mating-type locus (Gallagher et al. 2018; Torres-Garcia et al. 2020; Yaseen et al. 2022; Fellas et al. 2025). Such ectopic ’heterochromatin islands’ reduce the expression of embedded or nearby genes.

Directed screening for unstable caffeine resistance in wild-type fission yeast resulted in the isolation of epimutants that exhibited distinct H3K9me-heterochromatin islands at different chromosomal locations (Torres-Garcia et al. 2020; Yaseen et al. 2022; Fellas et al. 2025). Decreased expression of genes associated with such islands was shown to mediate caffeine resistance and characterized epimutants were also FLC resistant. In *S. pombe*, siRNA production is linked to Clr4 H3K9-methyltransferase recruitment and heterochromatin domain integrity (Martienssen and Moazed 2015; Allshire and Madhani 2018; Allshire and Ekwall 2015; Grewal 2023). Unique siRNAs, homologous to H3K9me heterochromatin island loci, were generated in caffeine-resistant epimutants and RNAi was shown to contribute to the resulting H3K9me heterochromatin-mediated resistance (Torres-Garcia et al. 2020). Similarly, unstable FK506 resistance in *Mucor circinelloides* isolates resulted from H3K9me-heterochromatin and/or homologous siRNA reducing expression of the *fkbA* gene (Son et al. 2026).

These observations suggested that heterochromatin-mediated epimutations could be responsible for the unexplained fluconazole heteroresistance observed in the euploid *C. neoformans* isolates within the Tanzanian patient cohort (Stone et al. 2019). Here, we demonstrate the presence of variable ectopic islands of H3K9 methylation in these unusual heteroresistant *C. neoformans* isolates. Both resistance and heterochromatin islands are lost upon prolonged growth in the absence of fluconazole and reappear upon drug re-exposure. Our analyses indicate that an alternative mechanism for development of heteroresistance, distinct from aneuploidy, can occur during *C. neoformans* infections which likely involves unstable heterochromatin-dependent repression of island-associated genes. Such an epigenetic route to FLC resistance has important implications for resistance characterization in cryptococcosis.

## Results

### Euploid *C. neoformans* heteroresistant clinical isolates exhibit ectopic islands of H3K9me2-coated chromatin

Heteroresistance has been associated with aneuploidy in clinical and environmental isolates of *C. neoformans* (Mondon et al. 1999; Yamazumi et al. 2003; Sionov et al. 2009, 2010; Yang et al. 2021). However, whole-genome sequencing (WGS) analysis of *C. neoformans var. grubii* samples obtained from twenty Tanzanian patients showed that, although all exhibited FLC heteroresistance, eight did not display aneuploidy (i.e. they were mainly euploid, containing one copy of each of the 14 chromosomes) and no strong candidates for causative genetic mutations were identified (Stone et al. 2019).

To determine how FLC heteroresistance might arise in these euploid isolates, we focused our analysis on four baseline (pre-treatment) isolates from this cohort (13R, 16R, 18R and 19R). These isolates were obtained from patients prior to FLC treatment but nevertheless formed FLC-resistant colonies on agar plates containing FLC (YPD+FLC; Fig. 1A). From the same cohort, isolate 11R had been shown to exhibit chromosome 1 aneuploidy; it served as an aneuploid control in our subsequent analyses. Whole genome sequencing confirmed the absence of aneuploidy in three of the four isolates analysed (Fig. 1B). However, isolate 19R contained a proportion of cells carrying more than one copy of chromosome 1. Prolonged growth of the 13R, 16R, 18R, 19R and 11R isolates for 8 to 24 days in the absence of FLC selection resulted in the loss, or substantial decrease, in the ability of each isolate to form FLC-resistant colonies compared to initial day 0 cells and the standard KN99 *C. neoformans var. grubii* reference isolate (Arras et al. 2017) (150 μM; Fig. 1C). Thus, as originally reported (Stone et al. 2019), all five tested isolates exhibited unstable resistance, characteristic of heteroresistance. Because isolate 19R exhibited some chromosome 1 aneuploidy (Fig. 1B) it was excluded from further analyses.

**Fig. 1.**
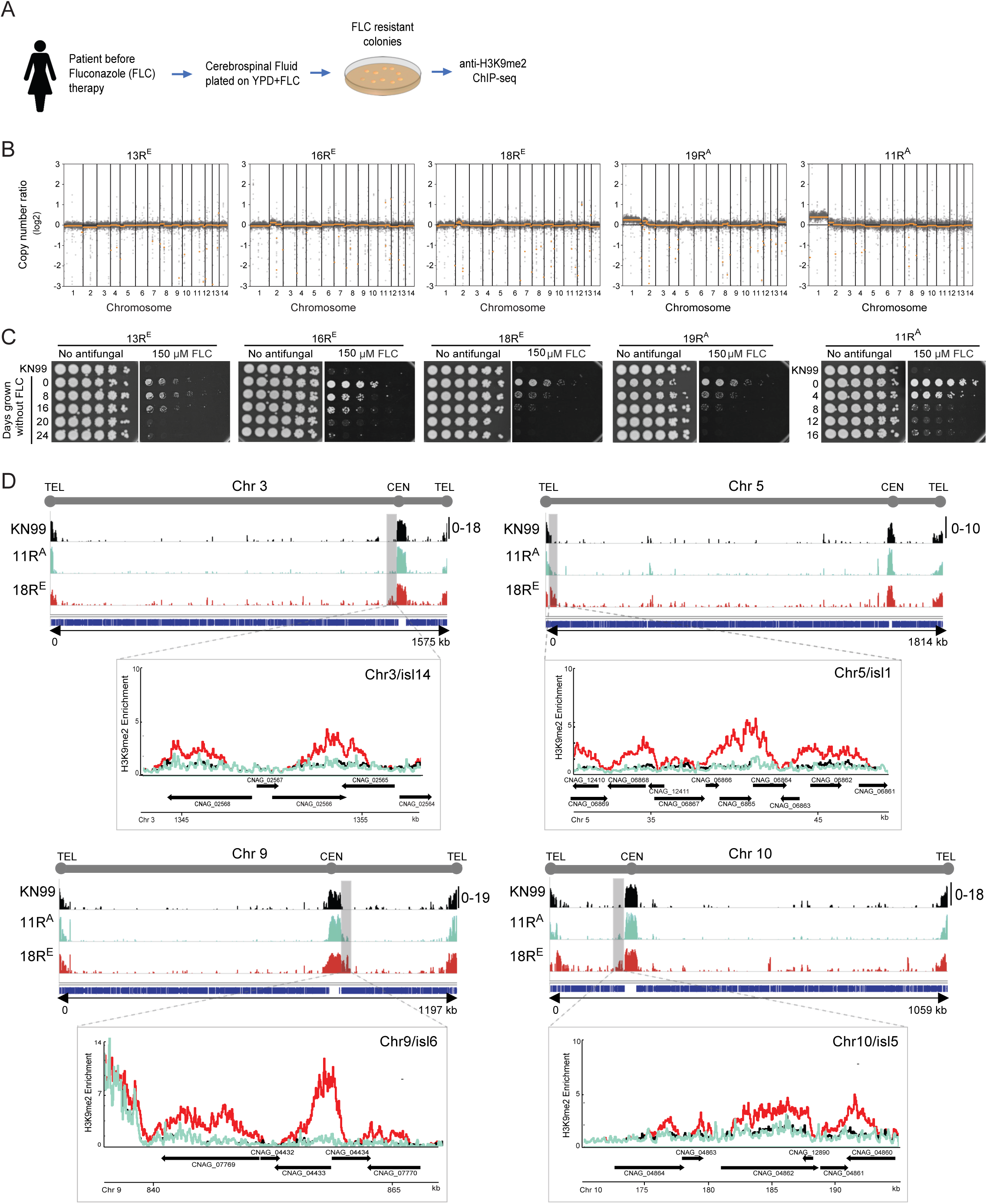
Euploid fluconazole-resistant clinical isolates exhibit ectopic H3K9me2 islands. (A) Schematic of experimental workflow. Cerebrospinal fluid samples from patients prior to fluconazole (FLC) treatment were plated on YPD+FLC to isolate resistant colonies (Stone et al. 2019), followed by anti-H3K9me2 ChIP-seq analysis. (B) Chromosome copy number variation (CNV) profiles of representative isolates (13R, 16R, 18R, 19R) and the aneuploid control 11R from genomic sequences. (C) Four-fold serial dilution growth assays on YPD plates with or without 150 μM FLC of isolates that had been propagated without FLC for the number of days indicated on left. KN99 is the reference wild-type control strain used. Plates were photographed after 4 days. (D) H3K9me2 ChIP-seq profiles of isolates indicated on left, across chromosomes 3, 5, 9, and 10 and enlarged view of across islands Chr3/isl14, Chr5/isl1, Chr9/isl6 and Chr10/isl5.

H3K9me heterochromatin-dependent epimutations have been shown to confer unstable resistance in other fungi by reducing the expression of associated genes (Torres-Garcia et al. 2020; Son et al. 2026). We reasoned that the fluconazole heteroresistance phenotype in the euploid isolates 13R, 16R and 18R might result from formation of heterochromatin islands at unusual chromosomal locations. *C. neoformans* exhibits both HP1/H3K9 methylation (H3K9me) and Polycomb/H3K27 methylation (H3K27me) heterochromatin systems (Dumesic et al. 2015).

To determine if either H3K9me- or H3K27me-dependent heterochromatin had accumulated at novel locations in the three euploid clinical isolates 13R, 16R and 18R, anti-H3K9 dimethylation (H3K9me2) and anti-H3K27 trimethylation (H3K27me3) ChIP-seq was performed and the resulting data compared to that for the KN99 reference strain (Fig. S1 and Table S1). In total, 122 H3K9me2 islands encompassing 359 genes were detected in these three clinical samples that were not present in the KN99 reference strain (Table S1). The overall distribution of H3K27me3 appeared similar to that of KN99 cells (e.g. 18R, Fig. S1, S2 and Table S1). Limitations on the amount of available anti-H3K27me3 precluded extensive investigation. Representative examples of H3K9me2 islands that were detected on chromosome 3, 5, 9 and 10 (Chr3/isl14, Chr5/isl1, Chr9/isl6 and Chr10/isl5) in isolate 18R (Fig. 1D), 13R and 16R (Fig. S3), but not KN99 cells or the aneuploid 11R^A^ clinical isolate, are shown. As it is well known that FLC heteroresistance often results from aneuploidy, the absence of these islands in aneuploid 11R^A^ suggested that the H3K9me2 islands present in non-aneuploid heteroresistant isolates may contribute to their FLC resistant phenotype.

#### Heteroresistant isolates lose fluconazole resistance and H3K9me2 islands following prolonged drug-free growth, and regain both when re-exposed to fluconazole

Unlike genetic mutants, RNAi-dependent and H3K9me heterochromatin-mediated epimutants have been shown to confer reversible drug resistance and lose resistance when grown in the absence of the selective compound employed (Calo et al. 2014, 2017; Torres-Garcia et al. 2020; Son et al. 2026; Fellas et al. 2025; Chang et al. 2019). Thus, the unstable characteristic of resistant epimutants resembles the heteroresistance phenotypes exhibited by several pathogenic fungi (Su et al. 2025).

To test if the presence of these islands is associated with fluconazole resistance, the 18R FLC-resistant isolate was propagated by repatching every 2 days on plates lacking FLC for 16 days; a non-clonal sample from each patch representing the population were frozen on days 0, 4, 8, 12 and 16 (Fig. 2A). Re-challenging cells from each non-clonal timepoint with 200 μM FLC showed that overall FLC resistance gradually declined during non-selective growth (Fig. 2B; compare day 0 with day 4, 8,12,16). Considerable loss of FLC resistance was apparent after growth for 16 days in the absence of FLC selection. We refer to this 16-day sample as 18R-revertant (18R-rev).

**Fig. 2.**
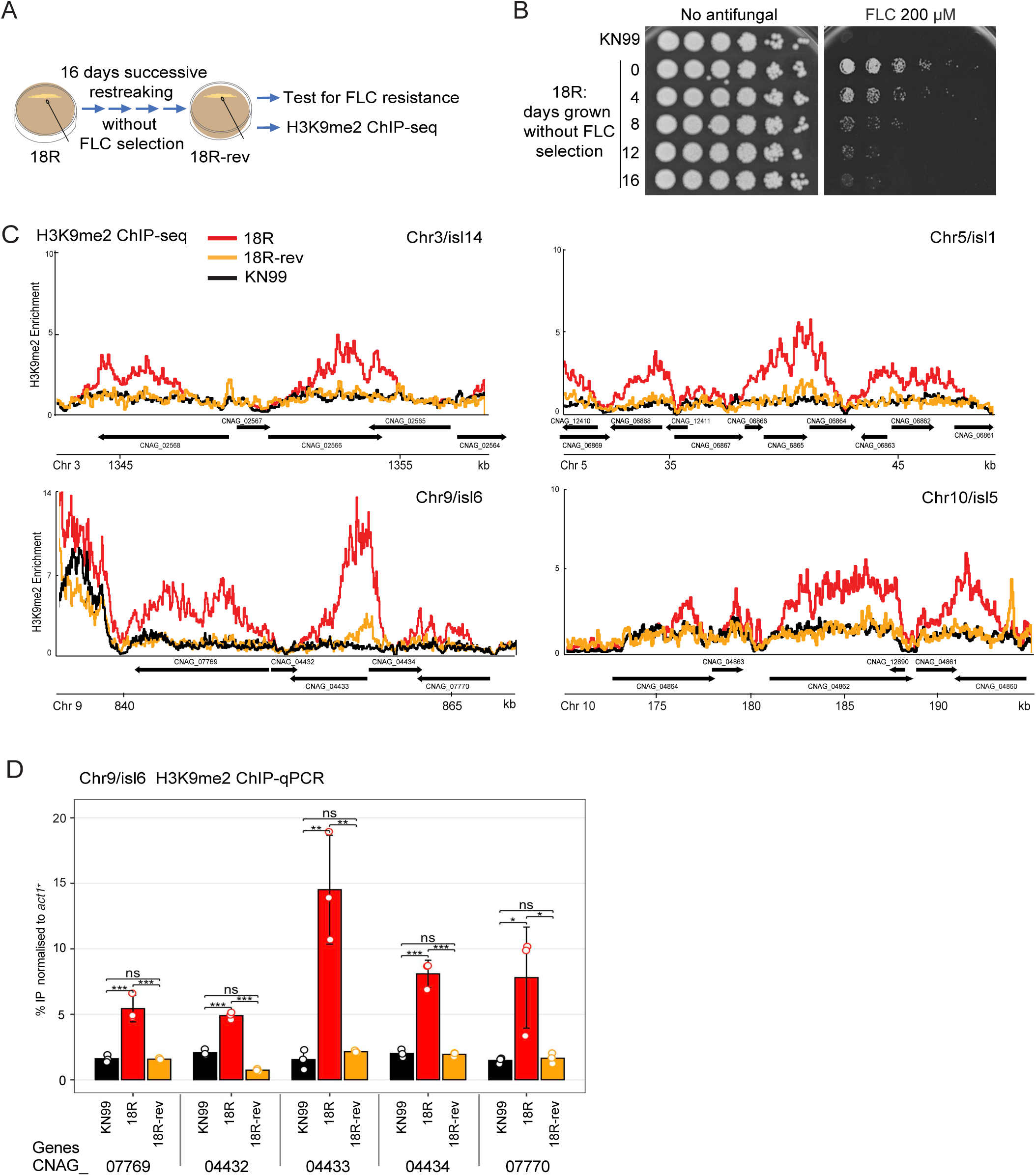
Loss of fluconazole resistance correlates with loss of H3K9me2 islands. (A) Experimental design for non-selective propagation of the 18R isolate to generate the sensitive derivative 18R-rev. (B) Serial dilution assays to assess loss of FLC resistance following non-selective (no FLC) propagation for the number of days indicated on left; YPD plates with or without 200 μM FLC. (C) H3K9me2 enrichment profiles across representative heterochromatin islands in KN99 control, 18R and 18R-rev cells (D) ChIP-qPCR validation of H3K9me2 levels at representative genes within Chr9/isl6. A one-way ANOVA test was applied (see Material and Methods): pvalue>0.05, ns; 0.01<pvalue <0.05,*; 0.001<pvalue<0.05, **; pvalue<0.001, ***.

ChIP-seq demonstrated that H3K9me2 levels at several H3K9me2-heterochromatin islands were reduced in 18R-rev compared to the parental 18R isolate, including Chr3/isl14, Chr5/isl1, Chr9/isl6 and Chr10/isl5 (Fig. 2C). A decrease in H3K9me2 levels on five specific genes within Chr9/isl6 was confirmed by ChIP-qPCR (Fig 2D) was confirmed by ChIP-qPCR (Fig 2D). Similarly, extended outgrowth of the 13R and 16R isolates in the absence of fluconazole resulted in 13R-rev and 16R-rev FLC-sensitive revertants that also exhibited decreased H3K9me2 levels at several islands (Fig. S3). In contrast, ChIP-seq revealed no notable gain or loss of H3K27me3 peaks between heteroresistant 18R cells and their largely FLC-sensitive 18R-rev derivatives (Fig. S2A-C). Moreover, the number of genes bearing H3K27me3 was almost identical in 18R and 18R-rev cells (Fig. S2C). Some genes exhibited both H3K9me2 and H3K27me3 (Fig. S2D). Because the clear variation in H3K9me2 distribution between resistant and revertant cells suggested a possible contribution to FLC resistance we focused subsequent analyses on H3K9 methylation.

If H3K9me2 heterochromatin islands contribute to the reversible FLC heteroresistant phenotype, then the re-exposure to drug should favor cells in the population with advantageous heterochromatin islands and thus select for their predominance. To explore this possibility, FLC-sensitive 18R-rev isolates that had lost H3K9me2 heterochromatin islands (Fig. 2B, C) were serially repatched on FLC-containing plates every 2 days for 8 days and two non-clonal populations isolated, 18R-revFLC1 and 18R-revFLC2, that exhibited greater growth on 150 μM FLC plates relative to the starting 18R-rev and KN99 cells (Fig. S4A and B). Increased H3K9me2 levels were detected at some islands in these 18R-revFLC cells relative to the initial FLC sensitive 18R-rev cells (Fig. S4C and D).

However, copy number variation (CNV) analysis of ChIP-seq input samples also revealed that 18R-revFLC2 cells were aneuploid for chromosome 1 (18R-revFLC2^A^), and 18R-revFLC1 apparently had a mixed population, with some cells displaying chromosome 4 aneuploidy (18R-revFLC1^A^) (Fig. S4E). Chromosome 4 disomy is also known to mediate FLC heteroresistance (Ngamskulrungroj et al. 2012). Aneuploidy and increased prevalence of H3K9me2 islands in FLC-resistant populations may either cooperate or act independently in distinct subpopulations to confer unstable resistance.

If aneuploidy and heterochromatin islands contribute to FLC resistance in 18R-revFLC1^A^ independently of each other, it should be possible to isolate euploid colonies that retain heterochromatin islands and associated FLC resistance. The 18R-revFLC1^A^ cell population that exhibited a low level of chromosome 4 aneuploidy was therefore streaked on plates containing FLC and five individual FLC-resistant colonies picked. These clonal isolates displayed different degrees of resistance to 200 μM FLC upon retesting in serial dilution assays (18R-revFLC1.1 to 1.5; Fig. 3A and B). ChIP-seq was performed on 18R-revFLC1.1-to-1.5. CNV analysis of input DNA revealed that two of the resulting single colony-derived isolates, 18R-revFLC1.4^A^ and 18R-revFLC1.5^A^, were aneuploid, whereas no aneuploidy was detected in 18R-revFLC1.1^E^, 18R-revFLC1.2^E^ or 18R-revFLC1.3^E^ cells (Fig. 3A). ChIP-seq showed that the euploid FLC resistant isolates 18R-revFLC1.1^E^, 18R-revFLC1.2^E^ and 18R-revFLC1.3^E^ exhibited increased H3K9me2 heterochromatin at various genomic loci including the islands Chr3/isl14, Chr5/isl1, Chr9/isl6 and Chr10/isl5 (Fig. 3C). In comparison, the aneuploid clonal isolates, 18R-revFLC1.4^A^ and 18R-revFLC1.5^A^, exhibited relatively lower levels of H3K9me2 at islands Chr3/isl14, Chr5/isl1, Chr9/isl6 and Chr10/isl5 compared to starting FLC resistant 18R cells (Fig. 3C). These observations suggest that, at least in 18R derivatives, FLC heteroresistance associated with aneuploidy can be separated from the presence of specific H3K9me2 heterochromatin islands. Aneuploidy, as well documented, provides one route to FLC resistance (Sionov et al. 2010; Su et al. 2025; Stone et al. 2019; Yang et al. 2021; Ngamskulrungroj et al. 2012). However, in euploid single colony isolates from the same cell population, FLC resistance is associated with the presence of islands of H3K9me2 heterochromatin. Although they may bolster heteroresistance in aneuploid cells, heterochromatin islands also appear sufficient to confer unstable FLC resistance alone.

**Fig. 3.**
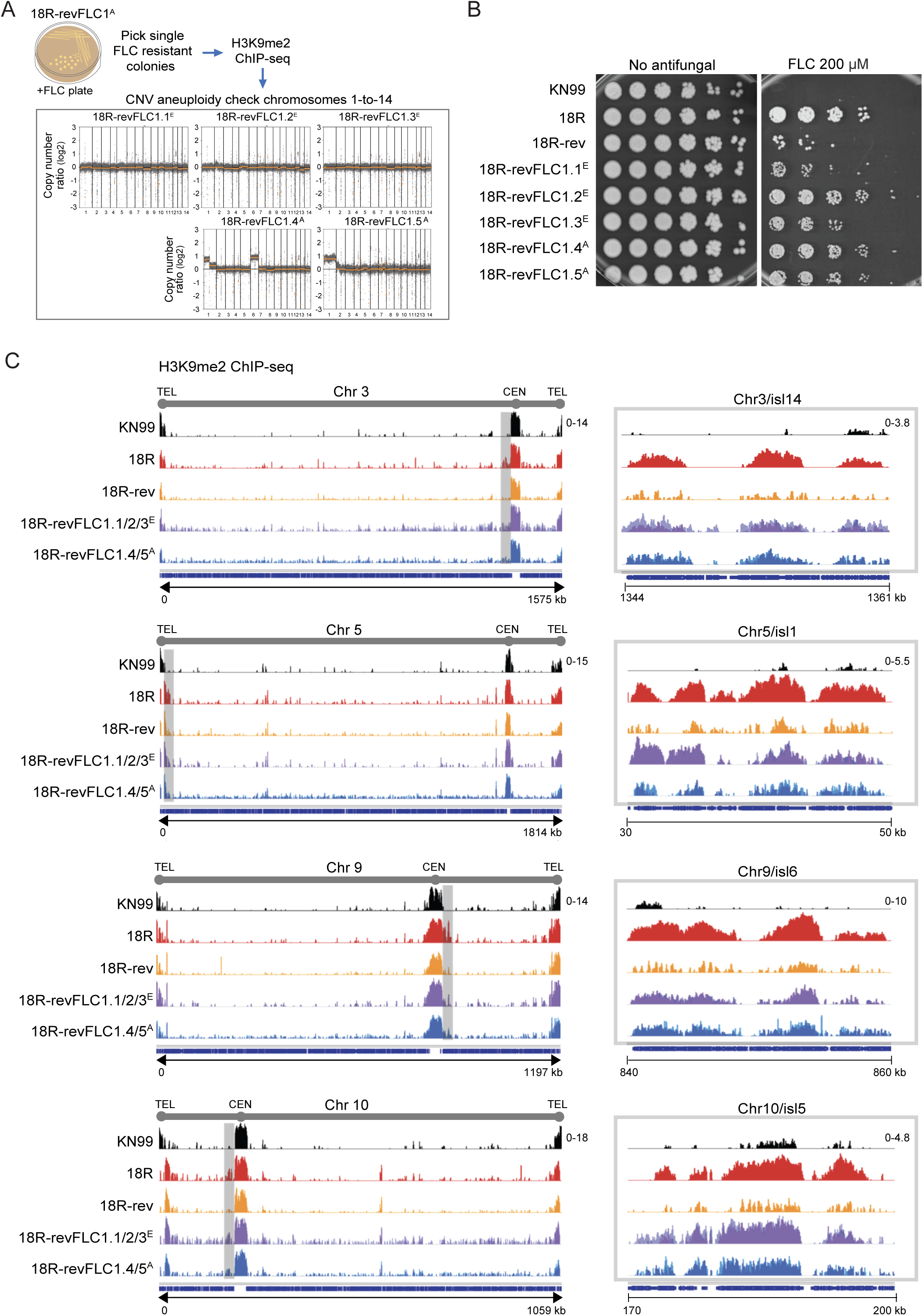
Fluconazole selection restores heterochromatin islands in resistant derivatives. (A) Experimental scheme for selection of FLC-resistant single colonies from the 18R-revFLC^A^ isolate which had a low level of chromosome 1 aneuploidy (top). CNV analysis from genome sequences from five single18R-revFLC^A^-derived FLC-resistant colonies (bottom). (B) Serial dilution growth assays of KN99 and 18R and 18R-rev cells and FLC-selected euploid and aneuploid derivatives on YPD plates with or without 200 μM FLC. (C) Genome browser views of H3K9me2 ChIP-seq profiles across chromosomes 3, 5, 9 and 10 to assess restoration of representative islands. Enlarged view of indicated islands in shaded regions is shown on right.

#### Fluconazole treatment of mice infected with *C. neoformans* 18R-rev allows recovery of heterochromatin islands and resistance

The analyses above demonstrate that H3K9me2 heterochromatin islands and FLC resistance co-vary in *C. neoformans* clinical isolates when grown in laboratory culture conditions. However, these isolates were originally obtained from infected patients. To investigate how H3K9me2 islands respond in a FLC-treated mammalian host, FLC sensitive 18R-rev cells were introduced into BALB/c mice by retro-orbital injection, which allows effective dissemination and a high frequency of brain colonisation (Coelho et al. 2019).

Infected mice were treated with FLC beginning 24 hours post-infection and every 24 hours thereafter. Seven days after infection, mouse brain (Mb) lysates were plated on YPD containing 150 μM FLC to recover FLC-resistant *C. neoformans* colonies. Colonies were retested for growth on 200 μM FLC and H3K9me2 distribution by ChIP-seq (Fig. 4A). ChIP-seq on cells from three FLC-resistant 18R-rev colonies recovered from mouse brain (18R-revFLC^Mb^1, 2 and 3) displayed H3K9me2 heterochromatin at Chr3/isl14, Chr5/isl1, Chr9/isl6 and Chr10/isl5 whereas no obvious chromosomal aneuploidy was detected (Fig. 4A and B). In contrast, FLC-resistant colonies recovered from the brains of mice infected with KN99 control cells displayed chromosome 1 aneuploidy without gain of H3K9me2 on island-associated genes (KN99FLC^Mb^1 and 2; Fig. S5). We conclude that FLC treatment during mouse infection with the 18R-rev isolate selects for the growth of cells with H3K9me2 islands that are associated with fluconazole heteroresistance.

**Fig. 4.**
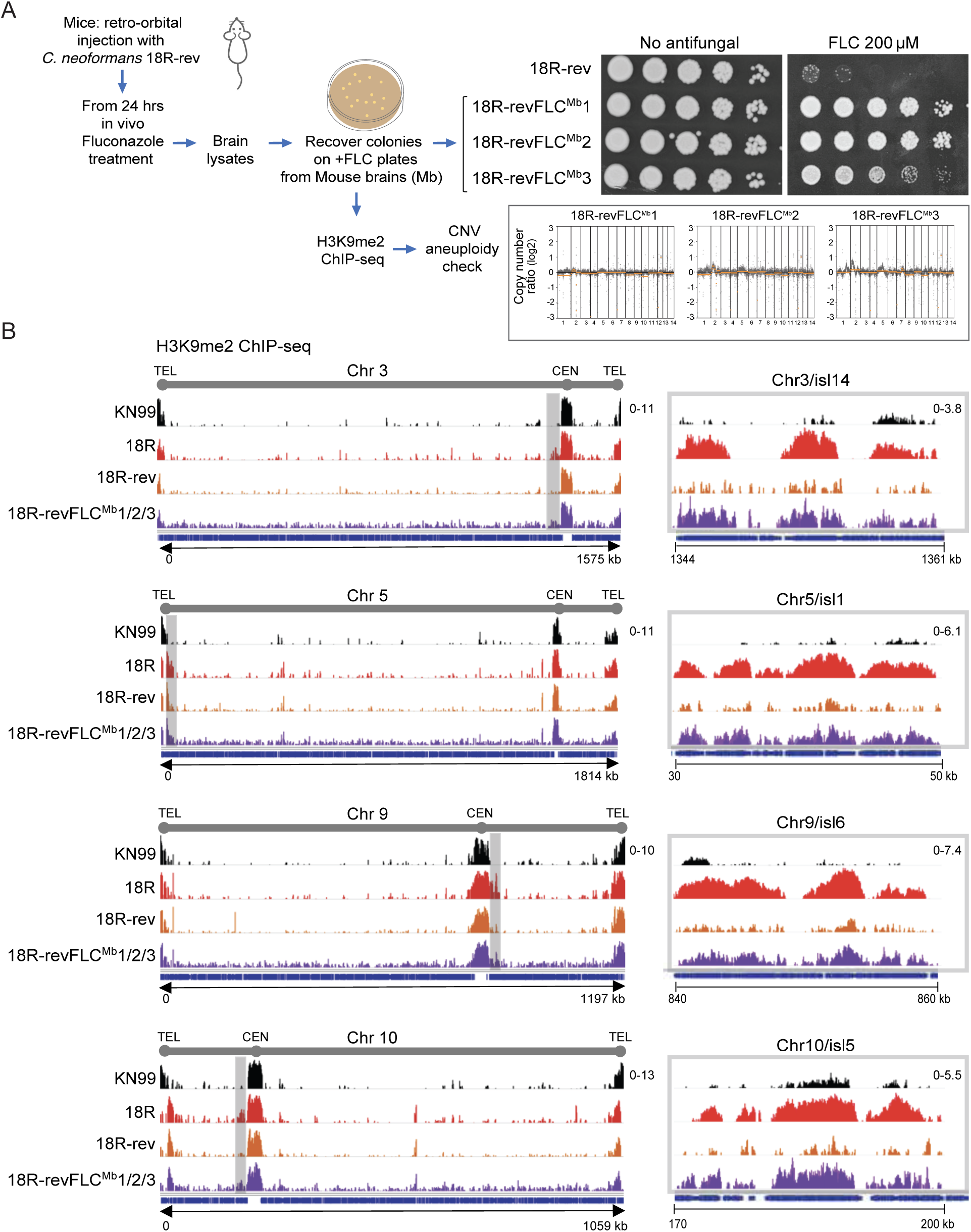
Fluconazole treatment *in vivo* restores heterochromatin islands. (A) Experimental scheme for recovery of FLC-resistant colonies from mouse brain following infection with 18R-rev, followed by fluconazole treatment (left). Serial dilution growth assay to assess FLC resistance of three isolates recovered from mouse brains (right). CNV analysis of the same isolates (bottom). (B) H3K9me2 ChIP-seq profiles to assess enrichment at representative islands in FLC-resistant euploid *C. neoformans* isolates recovered from brains of infected mice.

#### FLC-resistant isolates with variable H3K9me-islands exhibit subtle differences in associated gene expression relative to FLC-sensitive derivatives

Heatmap comparisons of the H3K9me2 ChIP-seq profiles of 13R-rev, 16R-rev, and 18R-rev FLC-sensitive derivatives confirmed a reduction in H3K9me2 relative to their FLC-resistant 13R, 16R and 18R progenitors (Fig. S6A, S7A). Thus, the loss of H3K9me2 from island-associated genes in these three euploid FLC-heteroresistant clinical isolates is associated with decreased FLC resistance.

Comparison of all *C. neoformans* genes carrying measurable H3K9me2 showed that 222 more genes exhibit measurable H3K9me2 in the original 18R FLC resistant isolate compared to its 18R-rev FLC sensitive derivative. H3K9me2 was again detected on over 449 genes in the 18R-revFLC1.1/2/3^E^ isolates that regained FLC resistance on plates, and 260 genes in the FLC resistant 18R-revFLC1/2/3^Mb^ isolates obtained from mouse brain tissue (Fig. S6B, S7D)

Gene Ontology (GO) terms of genes exhibiting higher H3K9me2 levels in FLC-resistant 18R cells relative to FLC-sensitive 18R-rev, revealed enrichment for genes encoding membrane components, transmembrane transport proteins, or transcription-related function (Fig. S6C). Membrane components and transmembrane proteins were also prevalent when GO term enrichment was assessed across genes exhibiting H3K9me2 within all three FLC-resistant isolates (13R, 16R and 18R) relative to their FLC-sensitive derivatives (Fig. S7B).

The co-variation of FLC-resistance with the presence of H3K9me2 islands suggested that the resistance phenotype might result from altered expression of genes located within or adjacent to these islands. RNA-seq was therefore conducted on 13R, 16R and 18R FLC^R^ cells and their 13R-rev, 16R-rev and 18R-rev FLC-sensitive derivatives. Within each isolate, a subset of genes displayed altered expression following non-selective outgrowth with an enrichment for gene ontology (GO) categories including membrane components, transmembrane transporters, and oxidative processes, known to be implicated in FLC response and broadly consistent with the GO analysis of H3K9me2-associated loci (Fig. S6D, S7C). However, no consistent transcriptional signature distinguishing FLC-resistance and FLC-sensitive states was detected across all three isolates. Given that these three isolates differ genetically, including isolate-specific SNPs and indels (Stone et al. 2019), it is likely that a combination of genetic background and the presence or absence of heterochromatin islands contribute to their overall resistance phenotype. Moreover, subtle and combinatorial differences in expression across multiple island-associated genes may contribute to resistance but remain difficult to detect using bulk RNA-seq in *C. neoformans*.

#### Most clusters of small RNAs are distinct from H3K9me2 islands

RNAi-dependent and heterochromatin-mediated epimutations have been shown to confer resistance to the protein phosphatase calcineurin inhibitor FK506 in the human fungal pathogen *Mucor circinelloides* (Calo et al. 2014; Son et al. 2026). In fission yeast, the integrity of H3K9me-dependent heterochromatin is reinforced by small RNA (sRNA) production both within constitutive heterochromatin domains and at heterochromatin island dependent epimutations (Allshire and Ekwall 2015; Martienssen and Moazed 2015; Allshire and Madhani 2018; Torres-Garcia et al. 2020; Grewal 2023). To determine if analogous sRNAs are produced from interstitial chromosomal loci associated with H3K9me2 islands in *C. neoformans,* sRNAs were extracted and sequenced from the 13R, 16R, 18R and 19R FLC-resistant clinical samples, their sensitive derivatives (13R-rev, 16R-rev, 18R-rev and 19R-rev) and KN99 cells (Fig. S8). Compared to H3K9me islands, fewer sRNA clusters were detected in the clinical isolates (an average of 358 H3K9me2 peaks versus 49 sRNA peaks for all isolates) that were not identified in the KN99 control. The location of sRNA clusters was variable across the four strains analysed and distinct from detected H3K9me2 peaks.

Many loci exhibiting sRNA clusters (35-42%) were common between KN99 cells and the 13R/13R-rev, 16R/16R-rev, 18R/18R-rev and 19R/19R-rev clinical isolates (Fig. S8C). The majority of these sRNA clusters (63-72%) were present in both the FLC-resistant (R) and matching sensitive (R-rev) derivatives; however, some specific clusters were only detected in the resistant (13-28%) or sensitive derivatives (9-16%) (Fig. S8C). Interestingly, one cluster of sRNAs mapped to the non-essential CNAG_06863 gene within H3K9me2 Ch5/isl1 in 18R but not 18R-rev cells (Fig. S8A and B). However, our RNA-seq analyses indicates that the presence of these siRNAs does alter CNAG_06863 expression. Moreover, published mutant analyses are inconclusive with respect to the possibility that loss or reduced CNAG_06863 function might confer FLC resistance (Billmyre et al. 2025; Boucher et al. 2025), and homology searches provide no indication of its possible function. The significance of these variable sRNA clusters remains to be determined.

#### Genes associated with H3K9me2 islands affect fluconazole resistance

Increased drug efflux due to an increase in the copy number of the *AFR1* gene, encoding an ABC transporter, is a major mechanism associated with chromosome 1 disomy and fluconazole heteroresistance in *C. neoformans* (Sionov et al. 2010; Chang et al. 2018).

Altered efflux might also contribute to the observed unstable FLC resistance of 18R cells. Rhodamine 6G (R6G) efflux assays demonstrated that aneuploid 11R cells exhibited increased R6G efflux compared to their euploid FLC-sensitive 11R-rev derivatives confirming that resistance is linked to chromosome 1 aneuploidy in 11R cells. However, no significant difference in efflux competence was observed between FLC-resistant 18R cells and their FLC-sensitive 18R-rev derivatives (Fig. 5A, B). Thus, the changes in gene expression that confer resistance in 18R cells must involve a mechanism that is distinct from increased efflux.

**Fig. 5.**
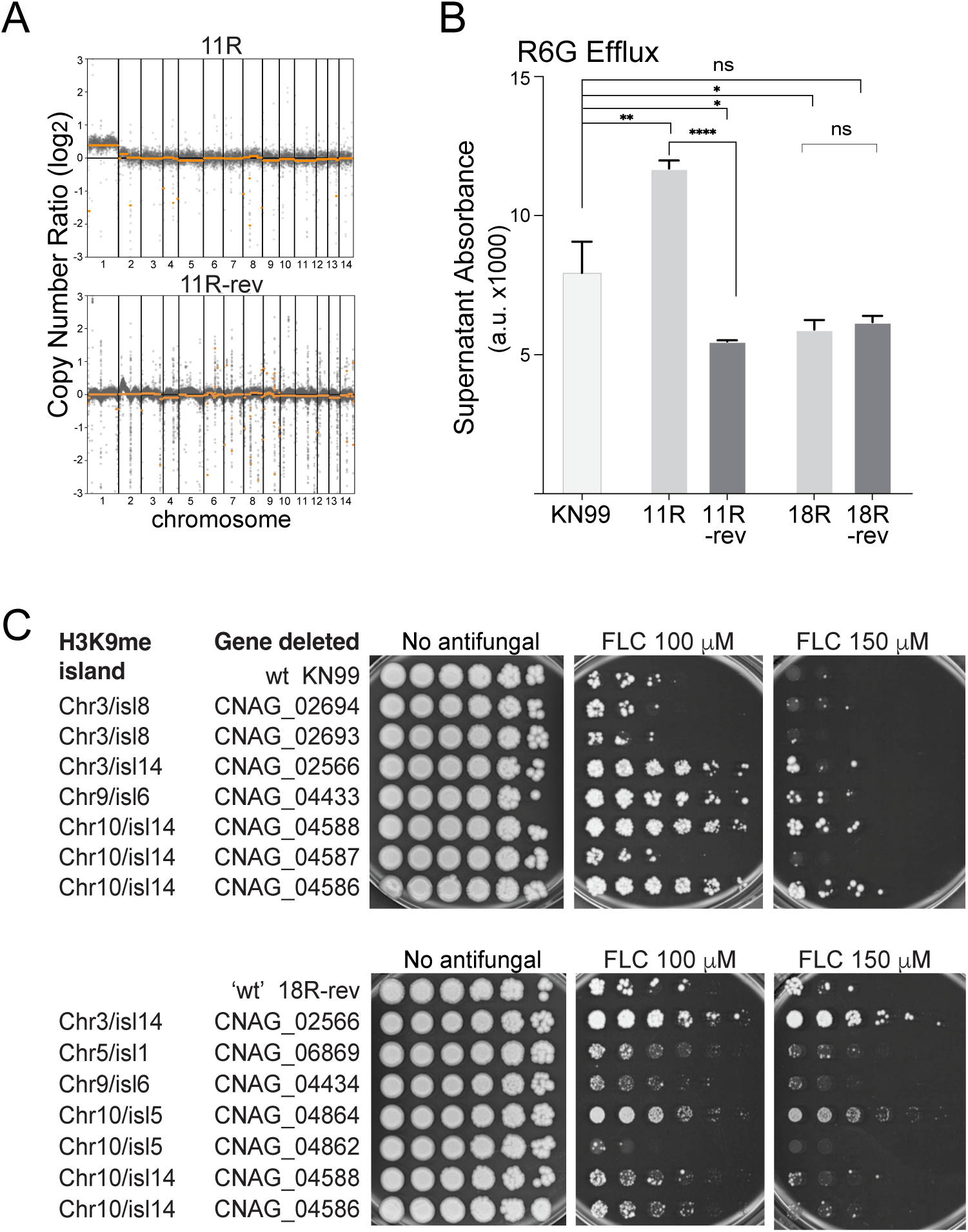
Island-associated genes influence fluconazole resistance. (A) CNV profiles of aneuploid FLC resistant isolate 11R and a revertant FLC sensitive derivative 11R-rev. (B) Rhodamine-6G efflux assay. Cells of the indicated strains were pre-loaded with Rhodamine-6G and glucose-induced efflux was measured by fluorescence after 1 hr. Three technical replicates were performed for each sample; standard deviation with *p* values determined by two-tailed Student’s t-test (p-value >0.05, ns; p-value <0.05, *; p-value<0.005, **; p-value<0.0001, **** GraphPad Prism) (C) Serial dilution growth assays of *C. neoformans* bearing deletions of the indicated genes that are associated with the specific heterochromatin islands as indicated. Gene deletions were obtained or generated in either the KN99 reference or 18R-rev backgrounds. Four-fold serial dilutions were performed, and cells were spotted on YPD plates containing the 100 or 150 μM Fluconazole (FLC).

H3K9me2 islands Chr3/isl14, Chr5/isl1, and Chr10/isl5 were consistently found to reappear in cell populations following growth in the presence of FLC (Figs. 3, 4; Fig. S7D). Both essential and non-essential genes are associated with these islands. Because heterochromatin islands are predicted to influence the expression of nearby genes, we determined whether non-essential island-associated genes contribute to FLC resistance.

KN99 cells harboring individual deletions of island-associated genes were obtained. Cells lacking (Δ) CNAG_02566 (Chr3/isl14), CNAG_04433 (Chr9/isl6), CNAG_04586 (Chr10/Isl14), or CNAG_04588 (Chr10/Isl14) displayed increased fluconazole resistance compared to wild-type KN99 cells on plates containing 100 or 150 μM FLC (Fig. 5C).

CNAG_02566 encodes the Fkh2 Forkhead-type transcription factor, CNAG_04433 a dual specificity serine/threonine DYRK protein kinase, CNAG_04586 the LIM-homeobox protein transcription factor Hob7, and CNAG_04588 the transcription factor Ert1.

As a more rigorous test, seven individual candidate genes located within or close to islands 2, 6, 7, 27, 28, and 29 were deleted in the FLC sensitive 18R-rev isolate. Cells in which either the CNAG_02566 (Fkh2 as above) or CNAG_04864 genes had been deleted exhibited greater fluconazole resistance on both 100 or 150 μM FLC plates relative to the parental 18R-rev derivative and other gene deletions (e.g. CNAG_04434 and CNAG_04862) isolated in the same background (Fig. 5C). CNAG_04864 encodes the Cir1 transcription factor involved in iron acquisition and regulates ergosterol biosynthesis. Both the Chr3/isl14 gene CNAG_02566/Fkh2 and the Chr10/isl5 gene CNAG_04864/Cir1 were previously implicated in resistance to azole-based antifungals (Jung et al. 2006, 2015). In addition, recent systematic screens identified defective CNAG_02566/Fkh2 or CNAG_04586/Hob7 function as increasing FLC resistance (Billmyre et al. 2025; Boucher et al. 2025).

The presence of genes linked to antifungal resistance within multiple heterochromatin islands suggests that the acquisition of heterochromatin at several chromosomal locations contributes to the observed unstable FLC resistance phenotype. The combined effect of multiple islands on the expression of genes located at distinct chromosomal loci likely contributes to the overall resistance phenotype. We conclude that modest alteration in the expression of numerous island-associated genes provides an alternative route to FLC heteroresistance distinct from the well-characterized mechanism of chromosomal aneuploidy.

## Discussion

Unstable heteroresistance in *C. neoformans* is typically associated with aneuploidy (Sionov et al. 2009), however, the identification of unstable FLC-resistant clinical isolates with euploid genomes suggested that other mechanisms might mediate heteroresistance (Stone et al. 2019). Here, we identify an epigenetic-based mechanism as an alternative route for heteroresistance in clinical isolates of *C. neoformans.* Ectopic heterochromatin islands enriched for H3K9me2 were detected at multiple genomic loci across three euploid clinical isolates from African patients with cryptococcal meningitis. Several of these H3K9me2 islands were lost following prolonged growth in the absence of FLC and were regained upon re-exposure to the drug, either *in vitro* or following FLC treatment of infected mice. Thus, the presence of specific H3K9me2 islands within a population correlates with a greater level of fluconazole resistance. In general, the presence of these variable H3K9me2 heterochromatin domains was associated with altered gene expression. However, no single causative gene was consistently identified across the three different euploid isolates examined. Instead, our analyses suggest that modest changes in the expression of several genes within each isolate contribute to the observed unstable FLC resistance phenotype.

Such changes may represent a novel type of epigenetically-mediated antifungal resistance where variable heterochromatin islands at several chromosomal locations contribute to the heteroresistant phenotype. Our analyses highlight the dynamic nature of heterochromatin as a potential adaptive mechanism for fungal persistence under antifungal selective pressure.

Four of the main H3K9me2 islands identified reside in close proximity to large domains of centromeric (Chr3_isl14, Chr9_isl6 and Chr10_isl5) or telomeric heterochromatin (Ch5/isl1). Such islands may not need to be established *de novo*; selection may allow growth of cells in which heterochromatin spreads from pre-existing sites to encompass genes whose altered expression confers a selective advantage (Allshire and Madhani 2018; Grewal 2023). This could include both the repression of genes that negatively influence survival under drug stress or altered expression of genes involved in processes such as detoxification, membrane transport or cellular stress responses (Fellas et al. 2025). Other islands identified in our study, such as Isl14 on chromosome 10, are distinct from centromeric and telomeric heterochromatin domains but they may also represent spreading from pre-existing nucleation sites to encompass more genes following FLC selection.

RNA-seq analysis did not identify specific candidate genes whose repression consistently explained resistance across the three euploid strains examined (13R/13R-rev, 16R/16R-rev, 18R/18R-rev; Fig. S6, S7). The intensity and extent of individual heterochromatin islands likely vary between individual cells in the population. Moreover, moderate alterations in the expression of genes within different islands may collectively contribute to resistance. Such incremental changes would be difficult to detect through transcriptome analyses. Previous studies in fission yeast demonstrated that even a 50% reduction in the expression of an essential mitochondrial gene is sufficient to confer resistance phenotypes (Torres-Garcia et al. 2020). Similarly, a 10-20% reduction in the expression of multiple genes affecting distinct pathways could generate resistance while avoiding the fitness costs associated with strong repression of any single gene. Functional analysis of candidate genes located within or adjacent to heterochromatin islands supported this interpretation. Deletion of several non-essential island-associated genes resulted in greater resistance to FLC (100 and 150 μM) in either KN99 cells (Chr3/isl14/CNAG_02566/Fkh2, Chr9/isl6/CNAG_04433/DYRK kinase, Chr10/isl14/CNAG_04587, Chr10/isl14/CNAG_04586/Hob7) or 18R-rev cells (Ch3/isl14/CNAG_02566/Fkh2 and Ch10/isl5/CNAG_04864/Cir1). These genes include transcriptional regulators and signalling proteins previously implicated in antifungal resistance pathways:

i. CNAG_02566/Fkh2 transcription factor regulates mitotic genes in both *S. cerevisiae* and *S. pombe* (Garg et al. 2015), and *S. pombe* Fkh2 associates with HDAC complexes and has been implicated heterochromatin spreading from nucleation sites (Greenstein et al. 2022).
ii. Deletion of the gene encoding CNAG_04864/Cir1 has been shown to increase the expression of ABC transporters AFR1, PDR1 and MDR1 that are known to be involved in xenobiotic efflux.
iii. CNAG_04588/Ert1 is a Zn(2)-C6 zinc cluster transcript factor. The *S. cerevisiae* ortholog acts as a transcriptional activator and repressor to regulate the switch from fermentation to respiration by acting on genes encoding proteins involved in gluconeogenesis and mitochondrial function (Gasmi et al. 2014).
iv. CNAG_04586/Hob7 is a general stress responsive transcription factor involved in differentiation. The *S. pombe* ortholog Phx1 is involved in shifting metabolism from respiration to fermentation in stationary phase thereby reducing ROS induced stress (Kim et al. 2014).

Heterochromatin-dependent epimutations that alter metabolism as a result of mitochondrial dysfunction confer drug resistance in *S. pombe* (Fellas et al. 2025), however, it remains to be determined if the *C. neoformans* Fkh2, Ert1 Hob1, and Phx1 proteins perform similar functions to their *S. cerevisiae* and *S. pombe* counterparts and how their loss increases resistance to fluconazole.

We suggest that different clinical *C. neoformans* isolates acquired distinct patterns of H3K9me-dependent heterochromatin that modulate the expression of different combinations of genes. Variations in their sRNA pools or H3K27me-dependent heterochromatin may also result in gene expression differences. The cumulative effect of the resulting modest changes in expression mediates the observed unstable FLC resistance phenotype. Thus, although single gene deletions can identify individual H3K9me island-associated genes that can contribute to resistance, a partial reduction in the expression of multiple genes would allow the additive effect of multiple hypomorphic heterochromatin-dependent epimutations to mediate resistance with minimal fitness costs. We suggest that such traits be termed ‘poly-epigenetic’.

The formation and persistence of heterochromatin islands may itself be influenced by genetic variation affecting chromatin regulatory pathways. For example, variants that increase the activity of an enzyme involved in promoting heterochromatin formation, such as histone deacetylases, H3K9/H3K27 methyltransferase, or that reduce the activity of factors opposing heterochromatin formation (histone deacetylases, histone demethylases), could favour island formation (Toda et al. 2026; Bhatt et al. 2026; Zofall et al. 2016; Wang et al. 2015). Conversely, isolates with increased rates of chromosome segregation errors may preferentially generate resistance through aneuploidy.

Heterochromatin-mediated heteroresistance likely operates alongside the well-established mechanism of chromosomal aneuploidy in *C. neoformans* (Sionov et al. 2009). Indeed, some islands were also detectable in aneuploid isolates 18R-revFLC1^A^ and 18R-revFLC2^A^. It is possible that aneuploidy and heterochromatin islands arise independently in separate subpopulations. Alternatively, epigenetically-mediated heteroresistance may precede and facilitate the emergence of chromosomal copy-number changes. One possibility is that heterochromatin islands provide an initial adaptive response that, under continued selective pressure, may be replaced or reinforced by perhaps more stable genetic alterations such as aneuploidy. The opposite is also possible where aneuploidy is replaced by epimutations.

Indeed, when *S. pombe* was challenged with caffeine, an advantageous heterochromatin-mediated epimutation was detected before gene copy number variation, and both were lost following growth in the absence of caffeine selection (Torres-Garcia et al. 2020). However, adaptation of fungi to external insults such as antifungal drugs is likely to be stochastic so that any cell in a population exhibiting a difference that confers an advantage –allowing them to continue to divide– will be selected. Such differences could initially be: decreased or increased expression of a specific gene; heterochromatin-, or RNAi-mediated epimutations (Calo et al. 2017, 2014; Son et al. 2026; Torres-Garcia et al. 2020); gain or loss of transposons (Priest et al. 2022; Gusa et al. 2020); gain or loss of prions (Harvey et al. 2017); whole or partial chromosomal aneuploidy (Sionov et al. 2010, 2009); or specific gene amplification (Chow et al. 2012). Each of these mechanisms could contribute to overall heterogeneity within a population and the unstable resistance or other phenotypes that emerge.

Resistance arising through epigenetic mechanisms present challenges because, unlike genetic mutations and changes in ploidy, the underlying changes are not readily detected by conventional genome sequencing approaches that can identify resistance-associated mutations or copy-number changes. The findings presented here therefore highlight the importance of considering epigenetic variation as a contributor to population heterogeneity when analysing antifungal resistance in clinical isolates. The operation of both genetic and epigenetic pathways in mediating antifungal resistance underscores the remarkable plasticity of fungal pathogens. Aneuploidy, epimutations and perhaps other adaptive mechanisms may operate simultaneously within a population, increasing the probability that at least some cells survive antifungal exposure. Understanding how these processes interact will be important for improving strategies to detect, monitor and ultimately counteract antifungal resistance in fungal pathogens.

## Materials and Methods

### Strains, media, and growth conditions

All *C. neoformans* strains used in this study are listed in Table S2. Growth conditions and selection of fluconazole-sensitive and resistant derivatives are as described in Supplementary Information.

#### Mouse infections

Mouse infection with *C. neoformans* isolates and their recovery from mouse tissues were as described in Supplementary Information.

#### ChIP, ChIP-seq, RNA isolation, RNA-seq, Small RNA Isolation, sRNA-seq, and Analysis

ChIP fixation and chromatin extraction, RNA isolation, small RNA isolation along with ChIP-seq, RNA-seq and sRNA-seq library construction methodology and analysis were performed as described in Supplementary Information.

#### Quantitative ChIP–qPCR (qChIP)

ChIP-qPCR assay were performed as detailed in Supplementary Information.

#### Copy Number Variation analysis

Details of the CNV analysis method are provided in Supplementary Information.

#### Rhodamine R6G efflux assays

Rhodamine 6G was used to measure glucose induced efflux as described in Supplementary Information.

## Data availability

RNA-Seq, sRNA-seq and ChIP-seq data generated in this study have been submitted to GEO under accession number GSE327864.

## Acknowledgements

We are grateful to Vishnu Priya Krishnan, Edward Wallace, and Elizabeth Bayne for helpful discussions and comments on the manuscript. We gratefully acknowledge Shaun Webb for assistance in the Bioinformatics Core of the Discovery Research Platform for Hidden Cell Biology and Martin Wear of the Edinburgh Protein Production Facility for facilitating incubator and flow hood use.

## Author contributions

R.Y., S.C., P.T., A.L.P., M.I.N.M., C.P.A., J.H., and R.C.A. designed experiments; RY, SC, PT, MINM, CPA, performed research; M.S., A.L.P, N.R.H.S., and T.B. methods and tools; R.Y., S.C., P.T., A.L.P., M.I.N.M., and C.P.A. analyzed data; R.Y., S.C., P.T., A.L.P., and R.C.A. wrote the paper.

