## Supplementary Information: Text, Figs S1-S8 + Legends for "Variable ectopic heterochromatin islands provide an alternative route to antifungal heteroresistance in *Cryptococcus neoformans*"

**PDF contains:**

Additional Supporting Text:

- Detailed Materials and Methods
- Supplementary References
- Supplemantary Figures S1 to S8 plus Legends

#### Detailed Materials and Methods

##### Strains, media, and growth conditions

All clinical, reference and laboratory generated *C. neoformans* strains used in this study are listed in Table S2. All strains were grown in standard media and culture conditions. Experiments were performed with logarithmic phase cells grown at 30°C for lab strains and heteroresistant clinical isolates, in Yeast Peptone Dextrose (YPD) medium, containing 1% Yeast Extract, 2% Peptone, 2% Dextrose. For serial dilution growth assays, equal amounts of starting cells were serially diluted four-fold in sterile water and spotted onto YPD only or FLC-modified YPD medium. Cells were grown at 30°C or 37°C for 5-7 days then photographed. For drug-modified YPD media, the following fluconazole (Sigma, PHR1160) concentrations were used: 100, 150 or 200 µM (equivalent to 30.63, 45.9 or 61.3 µg/ml). Deletion strains were generated using Cas9-mediated recombination as previously described (Huang et al. 2021), using oligonucleotides described in Table S3.

##### Clinical isolates sampling, generation of unstable resistant and revertant isolates

Following informed consent, cerebrospinal fluid (CSF) were collected from HIV-infected patients with a confirmed first episode of Cryptococcal meningitis before initiation of fluconazole therapy (Stone et al. 2019). The CSF samples were directly plated on YPD agar with fluconazole concentration ranging between 8–64 µg/ml for 72 hours at 30°C. Individual resistant colonies (isolates) that emerged were collected and stored on microbank beads at -80°C. Whole genome sequencing identified colonies from patient samples as *C neoformans var grubii*.

To determine unstable resistance, the *C. neoformans* resistant isolates, from -80°C storage was patched on a YPD agar plate without FLC and allowed to grow at 30°C. From this seed *C. neoformans* population (day 0), a single patch was made on fresh YPD agar without FLC and incubated at 30°C. After 2 days, a single patch of cells was again made on a fresh YPD agar without FLC. This serial repatching of cells was repeated every 2 days for 16 days. On days 4, 8,12 and 16, cells from repatched population were frozen in 50% glycerol stock and stored at -80°C. The 16 day non-resistant, revertant, isolates were further repatched every 2 days for another 8 days on YPD agar without FLC to obtain 24 day revertant isolates. All passaged and frozen isolates were rechallenged with fluconazole. The day 16 isolate is referred to as revertant isolate.

To obtain populations of resistant isolates 18R-revFLC1$^{A}$ and 2$^{A}$, two separate patches were made from frozen revertant isolate on unmodified YPD agar at 30°C. After 3 days of growth, the two were serially repatched every 2 days on YPD agar modified with 150 µM FLC for 8 days at 30°C. The two FLC resistant isolate patches obtained after 8 days were frozen in 50% glycerol stock and stored at -80°C.

To obtain populations of resistant colonies 18R-revFLC1.1$^{E}$, 2 $^{E}$ and 3$^{E}$ and 18R-revFLC1.4$^{A}$ and 5$^{A}$ (Fig. 3A), resistant isolate 18R-revFLC1$^{A}$ was streaked on YPD agar modified with 150 µM FLC and incubated at 30°C for 5 days. Five single colonies were selected and frozen at -at 80°C.

**Mouse infections**

KN99 and 18R-rev *C. neoformans* cells were collected from YPD culture by centrifugation, washed twice with sterile phosphate-buffered saline (PBS) and resuspended at 1x10$^{7}$ cell/ml in PBS. Five-week-old BALB/c mice were anesthetized with isoflurane and infected with 5×10$^{5}$ cells per mouse by retro-orbital injection. Mice received 12 mg/kg of fluconazole via intraperitoneal injection starting 24 hours post-infection and then every 24 hours thereafter. Mouse survival was monitored daily, and euthanasia was performed via $CO_2$ exposure upon reaching humane endpoints. For the fungal burden analysis, lungs and brains from infected mice were recovered and plated in YPD supplemented with 150 µM fluconazole to select for colonies.

**Chromatin immunoprecipitation (ChIP)**

ChIP experiments were performed essentially as previously described (Tong et al. 2019), using anti-H3K9me2 (5.1.1, gift from T. Urano) or H3K27me3 (gift from H. Madhani, UCSF).

A 50 ml *C. neoformans* culture was grown to log phase. 1% paraformaldehyde (Sigma) was added and incubated at room temperature for 15 minutes to fix the cells. 1/20$^{th}$ volume of 2.5 M glycine was then added to stop fixation. Cells were then washed twice with ice cold PBS and then resuspended in 650 µl of ice-cold lysis buffer (140 mM NaCl, 50 mM HEPESKOH pH 7.5, 1 mM EDTA, 1% Triton-X100, 0.1% sodium deoxycholate). 1 mM PMSF (Thermo Scientific) and protease inhibitors cocktail (Roche) was added to the lysis buffer. 0.5 mL acid washed glass beads was added. Ten bead beating cycles consisting of 1 minute each and 2 minutes resting on ice was employed to lyse *Cryptococcus neoformans* cells. Centrifugation at 6000 x g for 8 min was done to pellet the insoluble chromatin fraction and washed with 1 mL lysis buffer. The pellet was then resuspended in 300 µl lysis buffer containing 0.2% SDS. Chromatin obtained was then sheared using a Bioruptor (Diagenode) for 28 min, 30 sec ON/OFF cycles (HIGH setting) to obtain fragments of about 150 bp. After sonication, 900 µl of lysis buffer without SDS was added to the sample and samples clarified by centrifuging at 17,000 x g for 20 min. 15 µl of supernatant obtained was used as input control and 1100 µl was used for immunoprecipitation. 6 µl of anti-H3K9me2 monoclonal antibody (m5.1.1, a gift from Takeshi Urano) or anti-H3K27me3 was used. 75 µl of pre-washed protein G Dynabeads was then added to the 1100 µl sample and immunoprecipitation was performed on a rotating wheel overnight at 4°C. Samples were washed for 10 minutes in lysis buffer then subsequently, the following washes: twice 10 min in lysis buffer containing 0.5 M NaCl; 10 min in wash buffer (10 mM Tris-HCl pH 8, 0.25 M lithium chloride, 0.5% NP-40, 1 mM EDTA, 0.5% sodium deoxycholate) and 10 minutes in TE buffer (10 mM Tris-HCl pH 8, 1 mM EDTA). Washed beads was resuspended in 150 µl ChIP elution buffer (10 mM Tris pH 8.0, 300 mM NaCl, 5 mM EDTA, 1% SDS) and 15 µl input samples were resuspended in 135 µl ChIP elution buffer. Both input and IP samples were incubated at 65°C overnight with shaking to reverse crosslinks. 2 µl DNase-free RNase (Roche, 0.5 µg/µl) was added to all samples and incubated at 37°C with shaking for 1 hr and then with 20 µl of proteinase K (10 mg/ml) at 55°C with shaking for 2 hrs. Immunoprecipitated DNA was purified using Monarch® PCR & DNA Cleanup Kit (5 µg) (NEB #T1130).

**Quantitative ChIP–qPCR (qChIP)**

The quantitative ChIP (ChIP-qPCR) procedure was as previously described (Torres-Garcia et al. 2020), except DNA recovered as follows: 10% Chelex-100 resin (BioRad) was prepared and 100 µl added to each input and IP sample. To extract DNA, samples were boiled at 100°C for 12 min. Samples were then incubated at 55°C with 2.5 µl of proteinase K (10 mg/ml) added. Samples were boiled at 100°C for 10 min to inactivate Proteinase K. Samples were then recovered in fresh tubes.

qChIP was performed and analysed by real-time PCR using Lightcycler 480 SYBR Green (Roche) using oligonucleotides listed Table S3. All ChIP enrichments were calculated as % DNA immunoprecipitated for the gene of interest relative to its corresponding input samples and normalized to % DNA immunoprecipitated for *act1⁺* gene. Histograms represent the averages of three biological replicates. Error bars represent standard deviations.

A one way ANOVA test was applied to qPCR data (pvalue>0.05, ns; 0.01<pvalue <0.05 ,*; 0.001<pvalue<0.05, **; pvalue<0.001, ***). The results were plotted using R package ggplot2 (https://ggplot2.tidyverse.org/) (Wickham 2016)

**ChIP–seq library preparation and data analysis**

Illumina-compatible libraries for ChIP DNA libraries were constructed using NEXTflex-96 barcode adapters (Bioo Scientific) and Ampure XP beads (Beckman Coulter) as previously described (Tong et al. 2019). Pooled libraries were sequenced on NextSeq 2000 by 50-bp paired-end sequencing. Approximately 6–20 million 75-bp paired-end reads were produced for each sample. After demultiplexing, adapters were trimmed from raw reads using Cutadapt (Martin 2011). Adapter trimmed reads were aligned to the H99 *C. neoformans* (CNA3, GCA_000149245.3) reference genome using Burrows-Wheeler Alignment Tool (BWA) (Li and

Durbin 2010). The bam files obtained were processed using Samtools (v1.3.1) (Li et al. 2009) and Picard-tools (v2.1.0) (http://broadinstitute.github.io/picard) for sorting, marking and removing duplicates then indexing. Bigwig coverage files were generated using BamCoverage (deepTools v2.0) and IP to input ratios were calculated using BamCompare (deepTools v2.0) (Ramírez et al. 2016) in SES mode for normalisation (Diaz et al. 2012). Peak calling was done using MACS2 (Zhang et al. 2008) in PE mode and broad peak calling (broad-cutoff <0.001). Peaks that were significant enriched across all biological replicates, with at least one sample showing an enrichment over input data greater than 2-fold, were identified as histone marker-enriched regions. All peaks identified across samples were merged together, and peaks located within 500 bp of each other were merged into a single island. Islands were named according to the chromosome followed by the identified island number (Phanstiel et al. 2014). Integrative Genomics Viewer v2.8.13 was used to view and generate track plots of ChIP–seq data (Robinson et al. 2011). Homer annotation (Heinz et al. 2010), was performed to identify genes associated with peaks based on the closest TSS (Stajich et al. 2011).

**Total RNA extraction and RNA-seq library construction and analysis**

RNA was extracted from 10OD of culture and RNA was extracted with TRizol according to manufacture protocol with some modifications. In brief, 500µl TRizol was added to each pellet and incubated on ice for 5min followed by addition of Chlorofom (100µl) and further incubation at room temperature for 10min. Samples were centrifuged for 15 min at 4°C, aqueous phase transferred to a new tube and 1 volume of 100% ethanol added. The mixture was then transferred to RNA Clean & Concentrator columns (Zymo Research) and DNAse treatment performed by adding 42ul RNA wash buffer (Zymo Research), 5ul 10x DNase buffer, 1ul Turbo DNase (Thermo Fisher Scientific) to each column and incubate 37°C for 30min. RNA was quantified using Nanodrop and Qubit. PolyA RNA was extracted using 500 ng total RNA and NEB polyA selection module (New England Biolabs E7490) and the library was prepared with NEBNext Ultra™ II Directional RNA Library Prep Kit (New England Biolabs E7490) according to manufacturer's protocol. Assessment of quality of resulting libraries was performed using a Bioanalyzer High Sensitivity DNA Assay Chip and Bioanalyzer instrument (Agilent Technologies) and quantification was performed using Qubit fluorometer (Life Technologies). After pooling, libraries were sequenced using Illumina NextSeq 2000 P3 (1.2B), 100 Cycles kit (50 bp paired end).

Paired-end sequencing reads were aligned to *Cryptococcus neoformans* H99 reference genome (Genome and transcriptome annotations were download from FungiDB version 68) using STAR (Stajich et al. 2011; Dobin et al. 2012; Patro et al. 2017). Transcript abundance was quantified with Salmon (Patro et al. 2017). For differential expression analysis, raw count estimates were summarized at the gene level and analyzed in R with DEseq2 (Love et al.

2014). Genes with low abundance (<10 reads) across all samples were excluded to reduce instability from sparse counts. A cutoff of q-value =< 0.01 and absolute log2(FC) >=1 was used to identify DEGs.

**Copy number variation analysis**

Copy number variation (CNV) was determined using CNVkit in Whole-Genome Sequencing (-wgs) mode (Talevich et al. 2016). H99 ChIP-seq input bam files were used as reference and FungiDB version 68 gene annotation with average bin size 1598 bp was used.

**Gene Ontology (GO) enrichment analysis**

Gene Ontology (GO) enrichment analysis was performed using the R package clusterProfiler (Yu et al. 2012). The annotation database was download from FungiDB version 68 (/Users/ptong/genomes/CryptococcusH99/FungiDB-68_CneoformansH99_GO.gaf).
GO terms were tested using the enricher function with Benjamini–Hochberg correction for multiple testing and gene set size thresholds of greater than 5.

**Rhodamine R6G efflux assay**

Rhodamine 6G was used to measure glucose induced efflux as previously described (Stone et al. 2019). *C. neoformans* cells were grown overnight to logarithmic phase at 30°C. Overnight cells were grown in fresh YPD for 4 hrs to log phase at 30°C. Cells were transferred into 50 ml falcon tubes and washed twice by centrifuging 2 mins at 3,500 x g with 30 ml glucose-free PBS. Pellets were resuspend in 10 ml glucose-free PBS. Rhodamine 6G was added at a final concentration of 10 μM, vortexed and incubated at RT for 2 hours to allow dye uptake. Cells were centrifuged to pellet. Pellets were washed two times with glucose-free PBS to remove external Rh6G. After external Rh6G was removed, cells were resuspended in 5ml glucose-free PBS. An aliquot of 750 μl cell suspension was taken into microfuge tubes. 250 μl of glucose-free PBS or PBS with 8 mM glucose (to induce efflux activity) was added to each tube to make a final volume of 1 ml. Tubes were incubated at 30°C for 60 minutes then centrifuged at 3,500 x g for 2 mins. 200 μl of supernatant was transferred to a 96-well plate and R6G was measured with a Spectramax M5 Multimode plate reader (Molecular Devices, Wokingham), using SoftMax Pro/SpecMaxMT software. Fluorescence was measured at an excitation wavelength of 527 nm and an emission wavelength of 555 nm. Three technical replicates were performed for each sample and mean calculated; standard deviation with *p* values determined by two-tailed Student's t-test (GraphPad Prism).

**Data availability**

RNA-Seq, sRNA-seq and ChIP-seq data generated in this study have been submitted to GEO under accession number GSE327864.

**Supplementary Information References**


Diaz A, Park K, Lim DA, Song JS. 2012. Normalization, bias correction, and peak calling for ChIP-seq. *Stat Appl Genet Mol Biol* **11**: Article 9.

Dobin A, Davis CA, Schlesinger F, Drenkow J, Zaleski C, Jha S, Batut P, Chaisson M, Gingeras TR. 2012. STAR: ultrafast universal RNA-seq aligner. *Bioinformatics* **29**: 15–21.

Heinz S, Benner C, Spann N, Bertolino E, Lin YC, Laslo P, Cheng JX, Murre C, Singh H, Glass CK. 2010. Simple Combinations of Lineage-Determining Transcription Factors Prime cis-Regulatory Elements Required for Macrophage and B Cell Identities. *Mol Cell* **38**: 576–589.

Huang MY, Joshi MB, Boucher MJ, Lee S, Loza LC, Gaylord EA, Doering TL, Madhani HD. 2021. Short homology-directed repair using optimized Cas9 in the pathogen Cryptococcus neoformans enables rapid gene deletion and tagging. *Genetics*.

Li H, Durbin R. 2010. Fast and accurate long-read alignment with Burrows–Wheeler transform. *Bioinformatics* **26**: 589–595.

Li H, Handsaker B, Wysoker A, Fennell T, Ruan J, Homer N, Marth G, Abecasis G, Durbin R, Subgroup 1000 Genome Project Data Processing. 2009. The Sequence Alignment/Map format and SAMtools. *Bioinformatics* **25**: 2078–2079.

Love MI, Huber W, Anders S. 2014. Moderated estimation of fold change and dispersion for RNA-seq data with DESeq2. *Genome Biol* **15**: 550.

Martin M. 2011. Cutadapt removes adapter sequences from high-throughput sequencing reads. *EMBnetJ* **17**: 10–12.

Patro R, Duggal G, Love MI, Irizarry RA, Kingsford C. 2017. Salmon provides fast and bias-aware quantification of transcript expression. *Nat Methods* **14**: 417–419.

Phanstiel DH, Boyle AP, Araya CL, Snyder MP. 2014. Sushi.R: flexible, quantitative and integrative genomic visualizations for publication-quality multi-panel figures. *Bioinformatics* **30**: 2808–2810.

Ramírez F, Ryan DP, Grüning B, Bhardwaj V, Kilpert F, Richter AS, Heyne S, Dündar F, Manke T. 2016. deepTools2: a next generation web server for deep-sequencing data analysis. *Nucleic Acids Res* **44**: W160–W165.

Robinson JT, Thorvaldsdóttir H, Winckler W, Guttman M, Lander ES, Getz G, Mesirov JP. 2011. Integrative genomics viewer. *Nat Biotechnol* **29**: 24–26.

Stajich JE, Harris T, Brunk BP, Brestelli J, Fischer S, Harb OS, Kissinger JC, Li W, Nayak V, Pinney DF, et al. 2011. FungiDB: an integrated functional genomics database for fungi. *Nucleic acids Res* **40**: D675-81.

Stone NR, Rhodes J, Fisher MC, Mfinanga S, Kivuyo S, Rugemalila J, Segal ES, Needleman L, Molloy SF, Kwon-Chung J, et al. 2019. Dynamic ploidy changes drive fluconazole resistance in human cryptococcal meningitis. *J Clin Investig* **129**: 999–1014.

Talevich E, Shain AH, Botton T, Bastian BC. 2016. CNVkit: Genome-Wide Copy Number Detection and Visualization from Targeted DNA Sequencing. *PLoS Comput Biol* **12**: e1004873.

Tong P, Pidoux AL, Toda NRT, Ard R, Berger H, Shukla M, Torres-Garcia J, Müller CA, Nieduszynski CA, Allshire RC. 2019. Interspecies conservation of organisation and function between nonhomologous regional centromeres | Nature Communications. https://www.nature.com/articles/s41467-019-09824-4#disqus_thread.

Torres-Garcia S, Yaseen I, Shukla M, Audergon PNCB, White SA, Pidoux AL, Allshire RC. 2020. Epigenetic gene silencing by heterochromatin primes fungal resistance. *Nature* **410**: 120.

Wickham H. 2016. ggplot2, Elegant Graphics for Data Analysis. *R*.

Yu G, Wang L-G, Han Y, He Q-Y. 2012. clusterProfiler: an R Package for Comparing Biological Themes Among Gene Clusters. *OMICS: A J Integr Biol* **16**: 284–287.

Zhang Y, Liu T, Meyer CA, Eeckhoute J, Johnson DS, Bernstein BE, Nusbaum C, Myers RM, Brown M, Li W, et al. 2008. Model-based analysis of ChIP-Seq (MACS). *Genome Biol* **9**: R137.

### Supplementary Figure S1

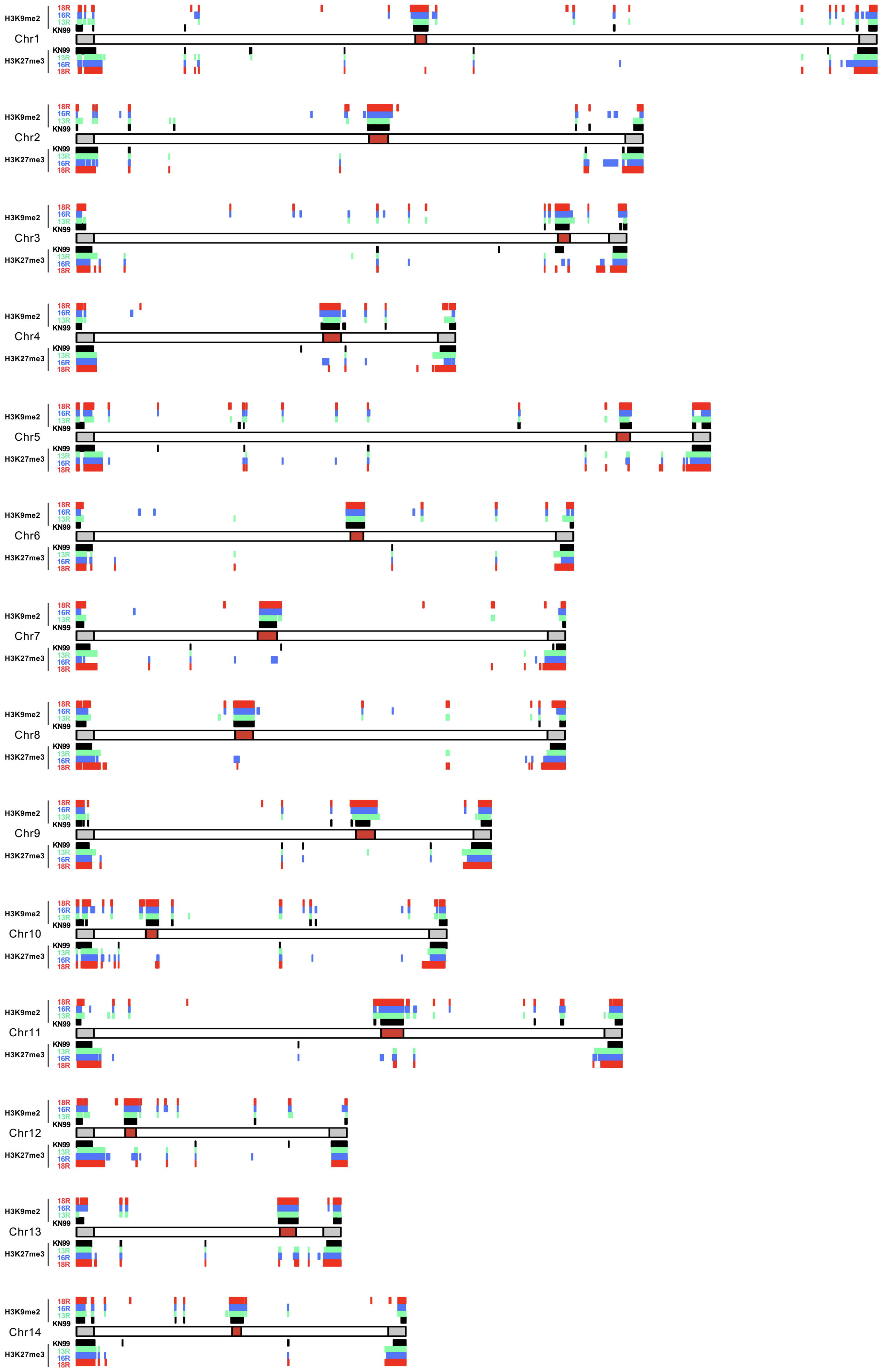

**Fig. S1. Genome-wide view of H3K9me2 and H3K27me3-enriched heterochromatin across *C. neoformans* chromosomes.**

A whole-genome view of chromatin islands enriched for H3K9me2 and H3K27me3 across all chromosomes. For each chromosome, H3K9me2-enriched regions are displayed above the chromosome ideogram, while H3K27me3-enriched regions are shown below. Enriched regions identified in the clinical strain KN99 are shown in black, whereas those fluconazole-resistant isolates are shown in green (13R), blue (16R), and red (18R). Chromosome ideograms are represented as horizontal bars, with centromeric regions indicated by red blocks and sub-telomeric regions (terminal 50 kb) highlighted in gray. The figure was generated using the karyoploteR package (https://bioconductor.org/packages/release/bioc/html/karyoploteR.html) .

### Supplementary Figure S2

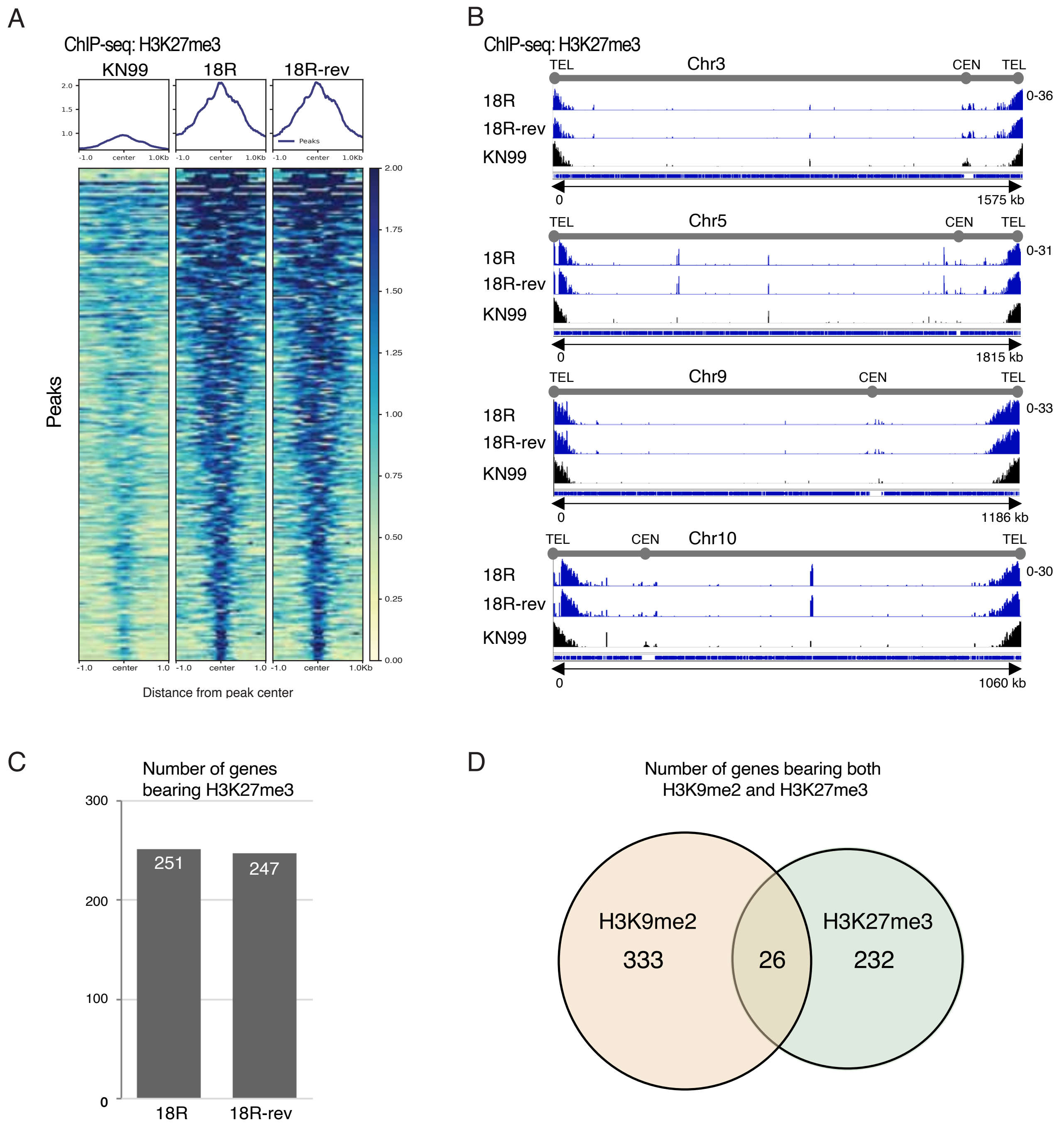

**Fig. S2**. **H3K27me3 distribution in FLC resistant and revertant isolates**

(A) Heatmap of H3K27me3 ChIP-seq enrichment peaks in KN99, 18R, and 18R-rev cells. Meta-profiles and ChIP-seq IP/Input normalised signal intensity across ±1 kb from center of peaks is shown.

(B) Genome browser views showing H3K27me3 enrichment across representative chromosomes in 18R, 18R-rev and KN99 cells.

(C) Histogram showing number of genes bearing H3K27me3 in 18R fluconazole-resistant cells and its 18R-rev fluconazole-sensitive derivative.

(D) Venn diagram showing number of genes exhibiting H3K9me2, H3K27me3 or both collectively specific to 13R, 16R, and 18R cells but not detected in KN99 cells.

### Supplementary Figure S3

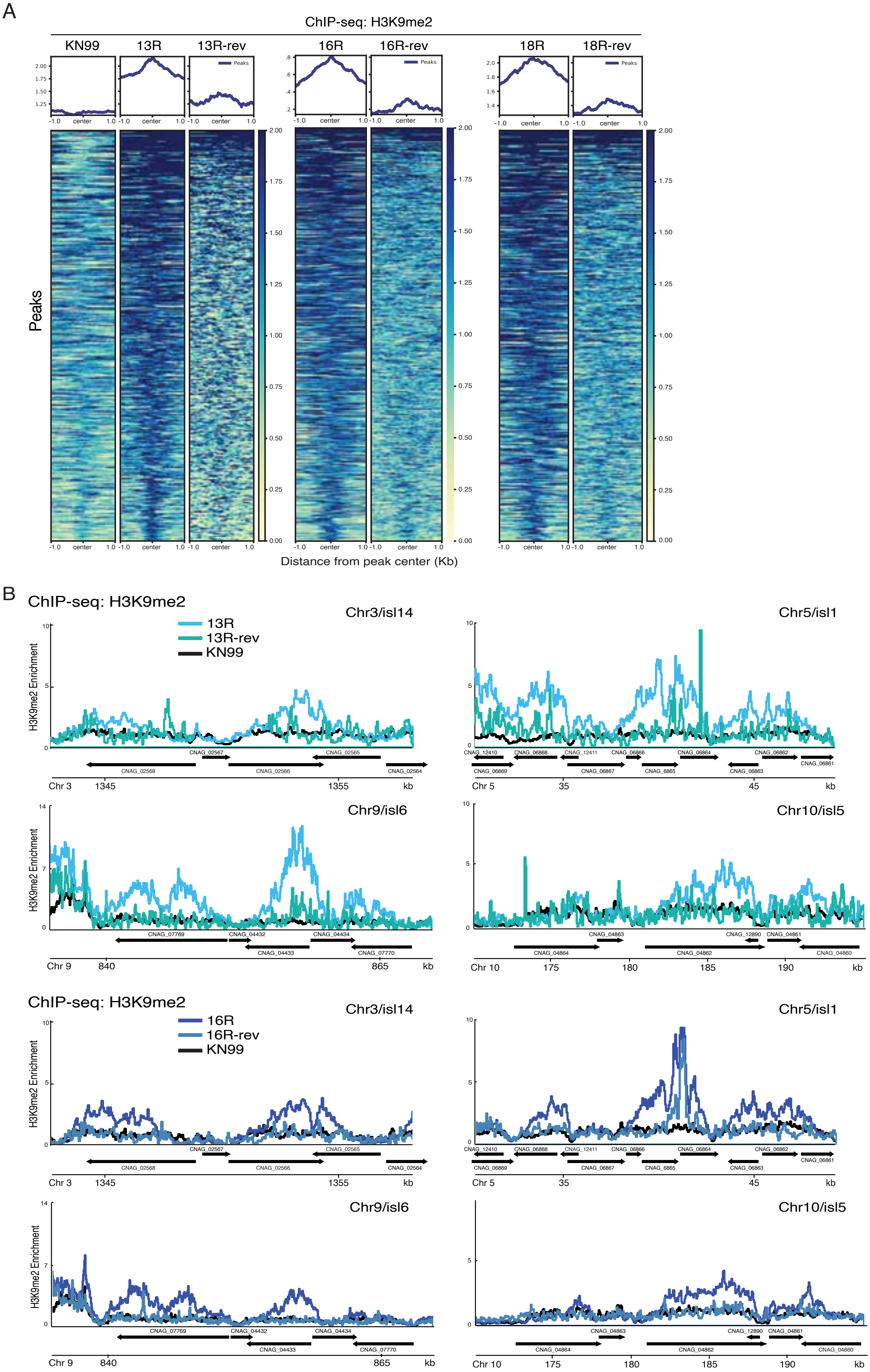

**Fig. S3**. **Loss of fluconazole resistance in 13R and 16R cells correlates with loss of H3K9me2 islands**

(A) Heatmaps of H3K9me2 ChIP-seq enrichment peaks in KN99, 13R, 13R-rev, 16R, 16R-rev, 18R and 18R-rev cells. Meta-profiles and ChIP-seq IP/Input normalised signal intensity across ±1 kb from center of peaks is shown.

(B) H3K9me2 enrichment profiles across the representative heterochromatin islands Chr3/isl14, Chr5/isl1, Chr9/isl6 and Chr10/isl5 in KN99, 13R, 13R-rev, 16R and 16R-rev cells.

#### Supplementary Figure S4

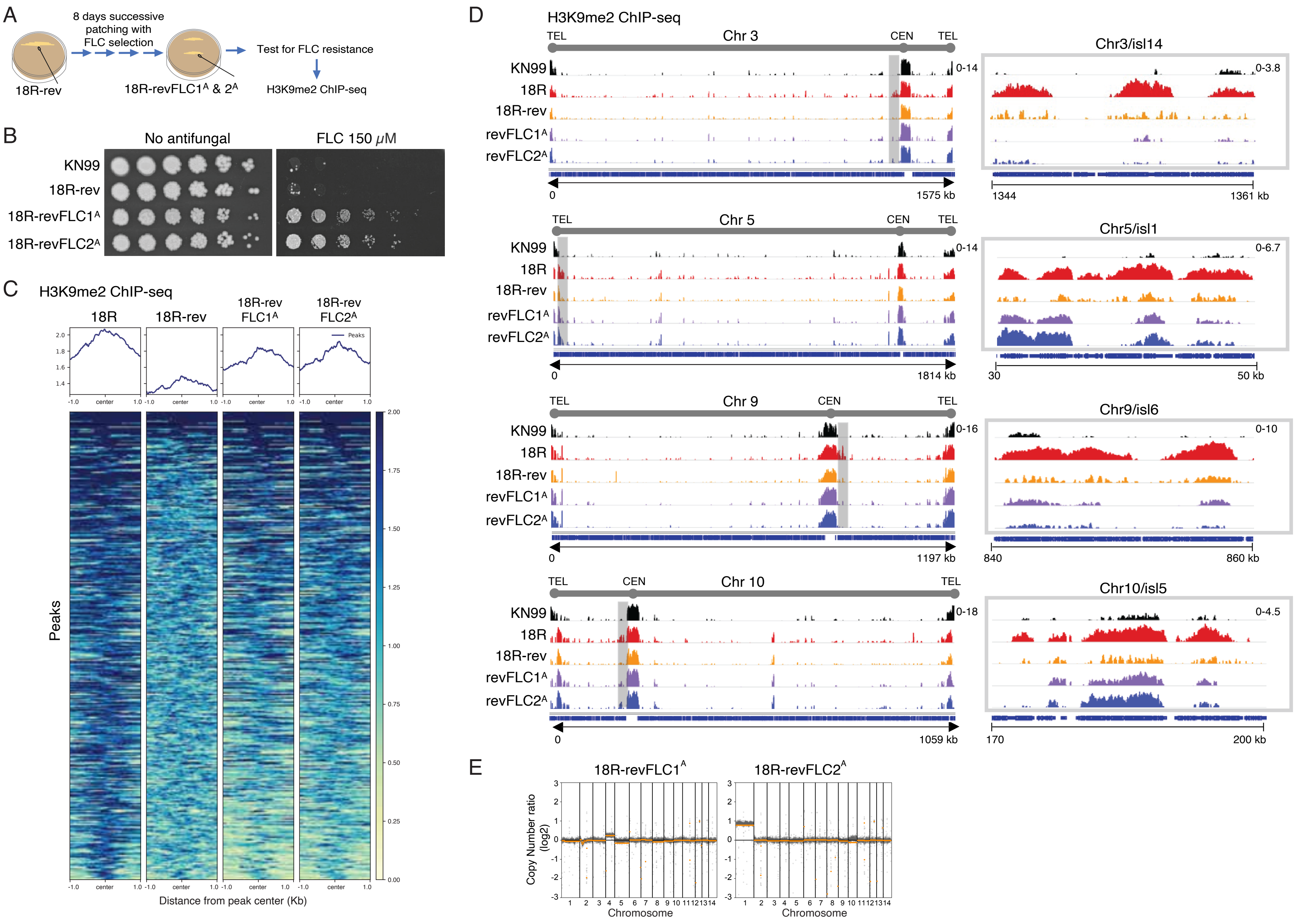

**Fig. S4**. **H3K9me2 profiles in aneuploid populations of fluconazole-selected 18R-rev cells**

(A) Experimental scheme for selection of FLC-resistant populations from 18R-rev cells.

(B) Serial dilution assays growth assays of KN99 and 18R-rev cells and FLC-selected populations (18-revFLC1$^{A}$ and 18-revFLC2$^{A}$) on plates with or without 150 μM FLC.

(C) Heatmap of H3K9me2 ChIP-seq enrichment peaks in 18R, 18R-rev and 18-revFLC1$^{A}$ and 2$^{A}$ populations. Meta-profiles and ChIP-seq IP/Input normalised signal intensity across ±1 kb from center of peaks is shown.

(D) Genome browser views of H3K9me2 ChIP-seq profiles across chromosomes 3, 5, 9 and 10 and enlarged view of islands Chr3/isl14, Chr5/isl1, Chr9/isl6 and Chr10/isl5 in KN99, 18R, 18R-rev and 18-revFLC1$^{A}$ and 18-revFLC2$^{A}$ populations.

(E) CNV analysis of 18-revFLC1$^{A}$ and 18-revFLC2$^{A}$ showing the presence of cells with an extra copy of chromosome 4 or 1, respectively.

### Fig. S4.

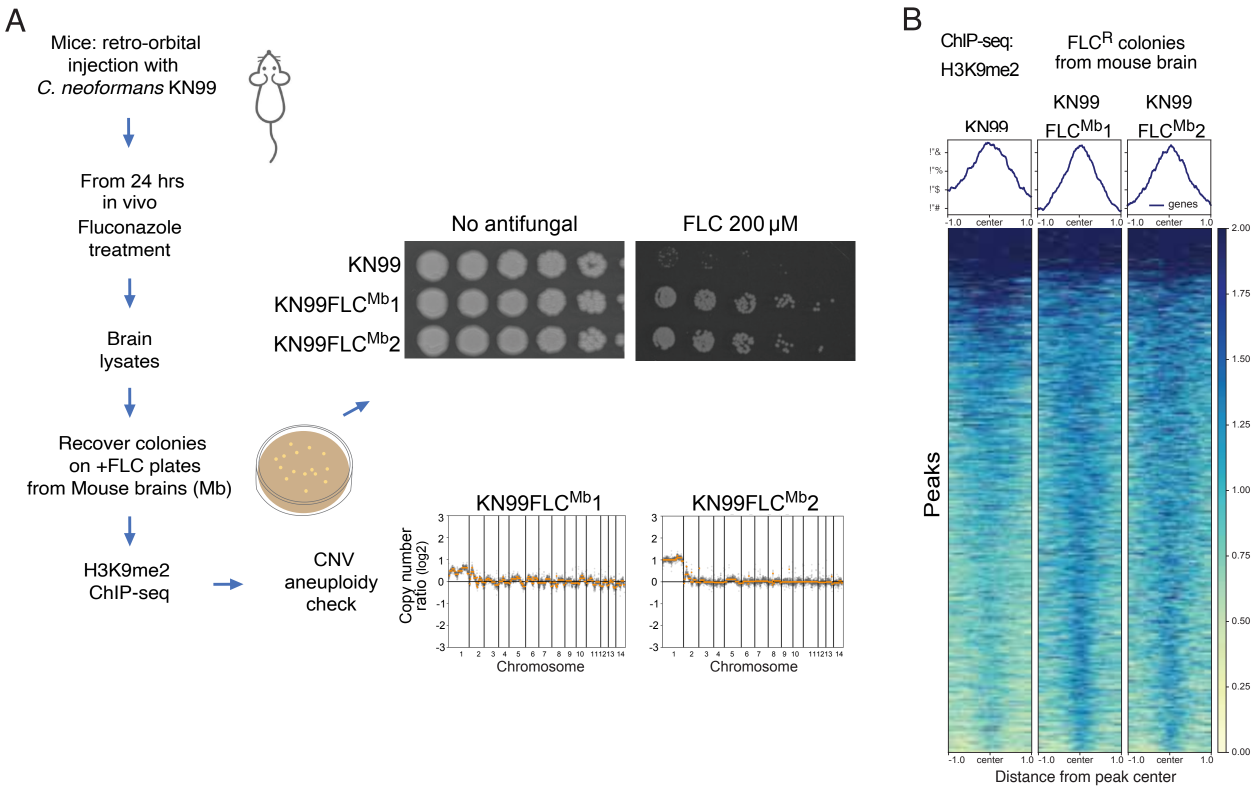
A
Mice: retro-orbital injection with C. neoformans KN99
From 24 hrs in vivo Fluconazole treatment
Brain lysates
Recover colonies on +FLC plates from Mouse brains (Mb)
H3K9me2 ChIP-seq
CNV aneuploidy check
No antifungal
FLC 200 µM
KN99
KN99FLCMb1
KN99FLCMb2
Copy number ratio (log2)
Chromosome
B
ChIP-seq: H3K9me2
FLCR colonies from mouse brain
KN99
KN99 FLCMb1
KN99 FLCMb2
genes
Peaks
-1.0 center 1.0
Distance from peak center
2.00
1.75
1.50
1.25
1.00
0.75
0.50
0.25
0.00

**Fig. S5. Fluconazole treatment of mice infected *in vivo* with KN99 cells results in chromosome 1 aneuploidy**

(A) Experimental scheme for recovery of FLC-resistant colonies from mouse brain following infection with KN99 and fluconazole treatment (left). Serial dilution growth assay to assess FLC resistance of two isolates from mouse brain (top right) and CNV analysis (bottom right). (B) Heatmaps of H3K9me2 ChIP-seq enrichment peaks in KN99 and fluconazole-resistant KN99 aneuploid isolates recovered from mouse brain. Meta-profiles and ChIP-seq IP/Input normalised signal intensity across ±1 kb from center of peaks is shown.

#### Fig. S6.

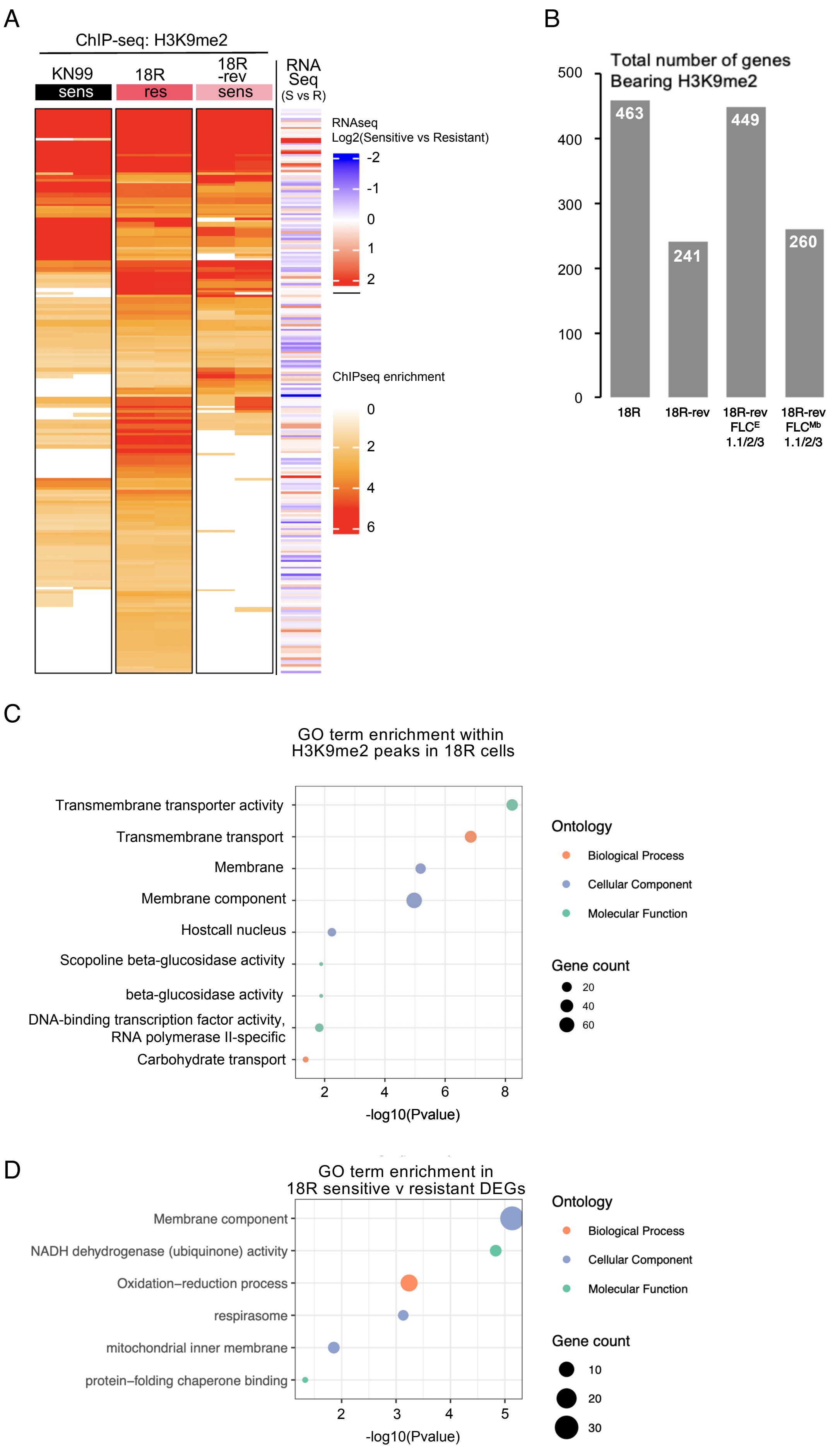
A
ChIP-seq: H3K9me2
KN99
sens
18R
res
18R -rev
sens
RNA Seq
(S vs R)
RNAseq
Log2(Sensitive vs Resistant)
-2
-1
0
1
2
ChIPseq enrichment
0
2
4
6
B
Total number of genes
Bearing H3K9me2
500
400
300
200
100
0
463
241
449
260
18R
18R-rev
18R-rev FLC^E 1.1/2/3
18R-rev FLC^Mb 1.1/2/3
C
GO term enrichment within
H3K9me2 peaks in 18R cells
Transmembrane transporter activity
Transmembrane transport
Membrane
Membrane component
Hostcall nucleus
Scopoline beta-glucosidase activity
beta-glucosidase activity
DNA-binding transcription factor activity,
RNA polymerase II-specific
Carbohydrate transport
2
4
6
8
-log10(Pvalue)
Ontology
Biological Process
Cellular Component
Molecular Function
Gene count
20
40
60
D
GO term enrichment in
18R sensitive v resistant DEGs
Membrane component
NADH dehydrogenase (ubiquinone) activity
Oxidation−reduction process
respirasome
mitochondrial inner membrane
protein−folding chaperone binding
2
3
4
5
-log10(Pvalue)
Ontology
Biological Process
Cellular Component
Molecular Function
Gene count
10
20
30

**Fig. S6. Transcriptional analyses of 18R versus 18R-rev cells relative to H3K9me2 island-associated genes**

(A) Heatmap of genes exhibiting H3K9me2 enrichment in KN99 reference strain and 18R-rev FLC-sensitive relative to 18R FLC-resistant cells along with $\log_2$ fold-change in the expression of genes associated with these H3K9me2 islands.

(B) Histogram showing number of genes exhibiting H3K9me2 in 18R FLC-resistant cells, 18R-rev FLC-sensitive cells, 18R-revFLC1.1/2/$3^{E}$ in vitro (from YPD plates) derived FLC-resistant cells or 18R-revFLC1.1/2/$3^{Mb}$ in vivo (from mouse brain) derived FLC-resistant cells

(C, D) Gene ontology (GO) enrichment analyses for H3K9me2-associated genes (C), and differentially expressed genes (D) in 18R-rev FLC-sensitive versus 18R FLC-resistant cells.

#### Supplementary Figure S7

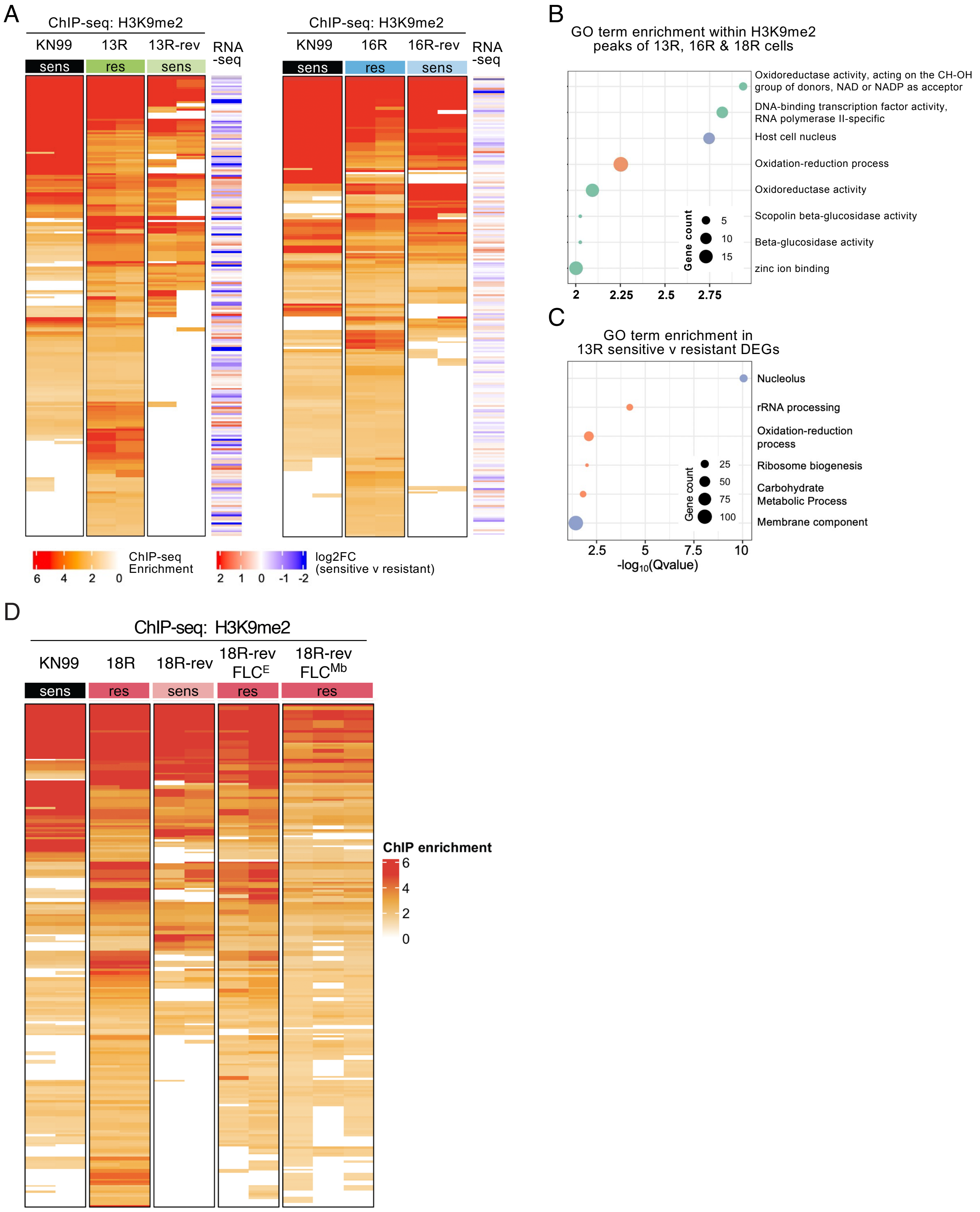

**Fig. S7. Transcriptional analyses of 13R and 16R versus 13R-rev and 16R-rev cells relative to H3K9me2 island-associated genes**

(A) Heatmap of genes exhibiting H3K9me2 enrichment in KN99 reference strain, 13R-rev FLC-sensitive relative to 13R FLC-resistant cells along with $log_2$ fold-change in the expression of genes associated with these H3K9me2 islands (left)
Heatmap of genes exhibiting H3K9me2 enrichment in KN99 reference strain, 16R-rev FLC-sensitive relative to 16R FLC-resistant cells along with $log_2$ fold-change in the expression of genes associated with these H3K9me2 islands (right).
(B) Gene ontology (GO) enrichment analyses for H3K9me2-associated genes shared by FLC-resistant 13R, 16R, and 18R cells.
(C) Gene ontology (GO) enrichment analyses of differentially expressed genes (DEGs) in 13R-rev FLC-sensitive versus 13R FLC-resistant cells.
(D) Heatmap of genes exhibiting H3K9me2 in KN99, 18R FLC-resistant, 18R-rev FLC-sensitive, 18R-revFLC1.1/2/3$^{E}$ *in vitro* (Euploid from YPD plates) derived FLC-resistant or 18R-revFLC1.1/2/3$^{Mb}$ *in vivo* (from Mouse brain) derived FLC-resistant cells

### Supplementary Figure S8

A

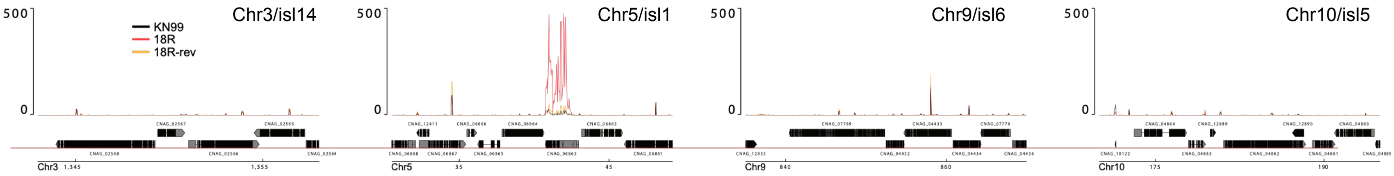


B

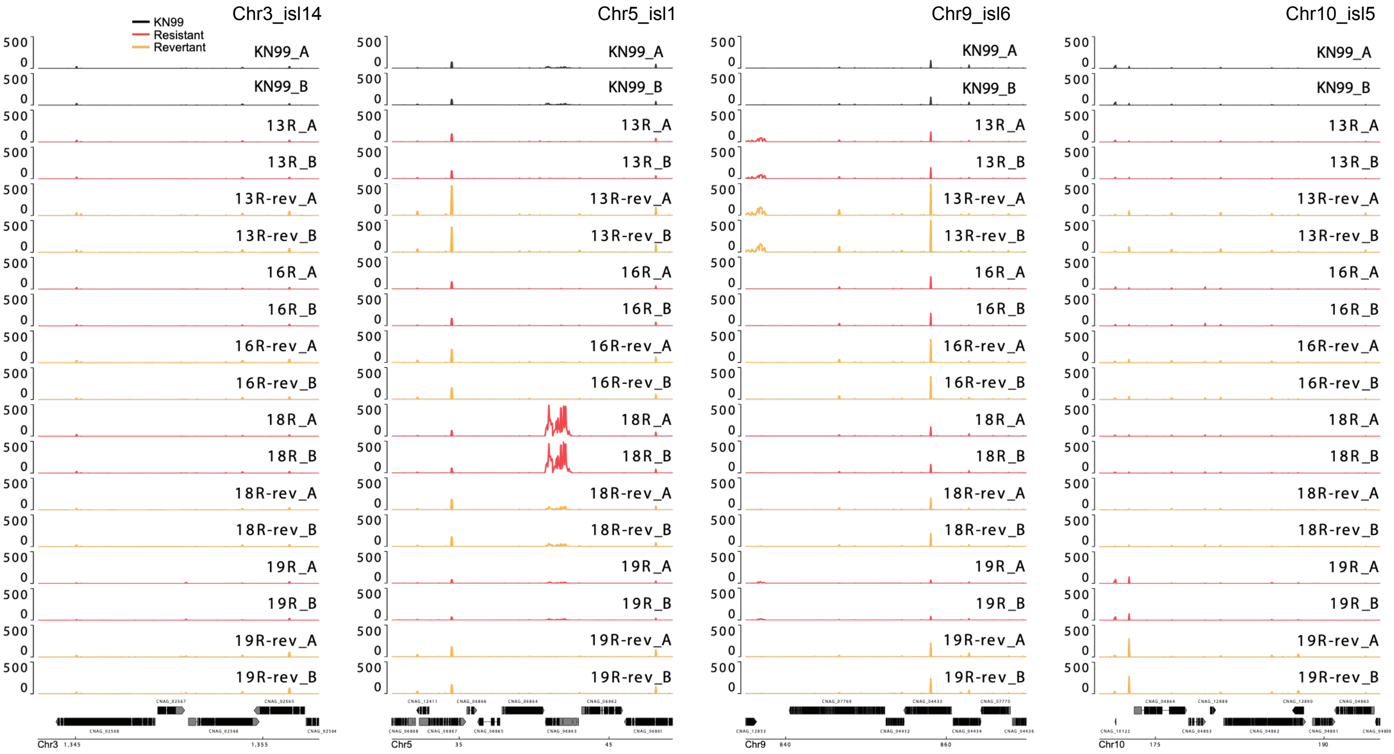


C

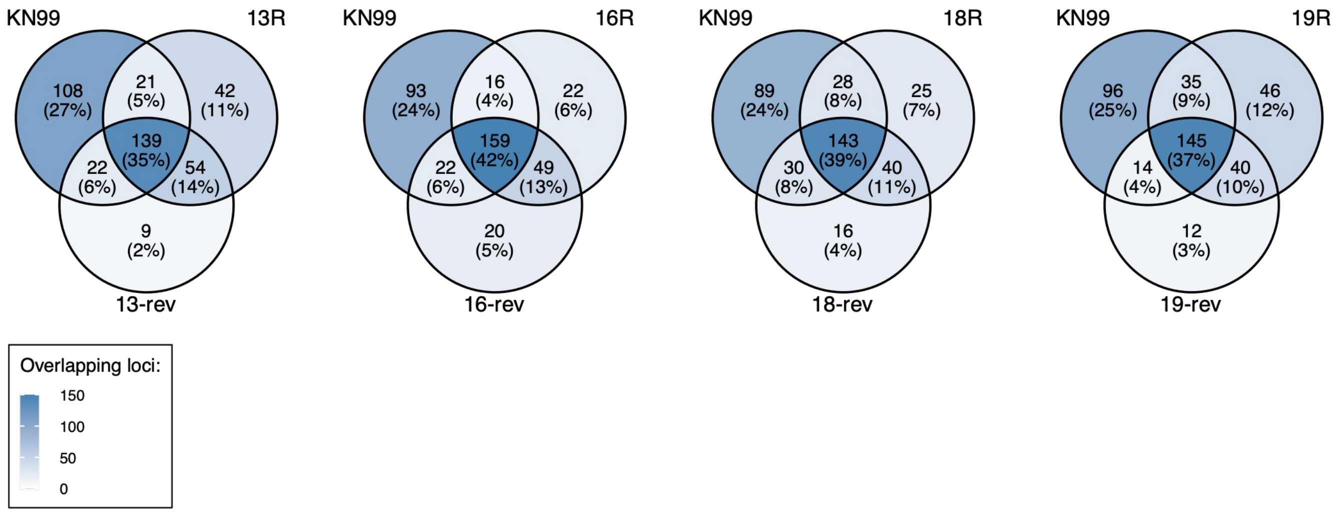

**Fig. S8**. **Comparison of small RNA levels in KN99, 13R, 13R-rev, 16R, 136R-rev, 18R, 18R-rev, 19R and 19R-rev cells**

(A-B) Genome browser views of sRNAs extracted from the indicated cells and mapped to regions of the genome containing the representative islands Chr3/isl14, Chr5/isl1, Chr9/isl6, Chr10/isl5.

(C) Venn diagrams showing the overlap in sRNA-associated genes between KN99, 13R, 13R-rev, 16R, 136R-rev, 18R, 18R-rev, 19R or 19R-rev cells
